# The role of pathogens in creating and maintaining host polymorphism

**DOI:** 10.1101/2025.02.13.638051

**Authors:** Edgard Djahoui, Nicolas Loeuille, Rudolf P. Rohr

## Abstract

Terrestrial and marine ecosystems exhibit remarkable species diversity and richness. Among the hypotheses put forward to explain this diversity —its origins and the forces sustaining it— one prominent hypothesis posits that pathogens could be pivotal in generating and maintaining species diversity within ecosystems. To explore this hypothesis, we analyzed a host-pathogen system, examining the specific conditions that might facilitate pathogen-driven diversification within the host population, with a focus on both the emergence and persistence of diversity. By applying a fecundity-transmission trade-off in which a non-evolving pathogen influences its evolving host, we observed that pathogens only infrequently drive host diversification, and only around intermediate trade-off conditions. Furthermore, while these rare diversification events result in stable coexistence, evolved diversity is low, consisting of two coexisting host morphs. Because the sole evolution of hosts in response to pathogens offers limited opportunity for diversification, we propose additional mechanisms that could foster higher levels of diversity within the system.

## 1 Introduction

In ecology and evolutionary biology, one of the central questions debated for more than a century is the origin and maintenance of the great species richness and diversity observed on Earth (Hutchinson, 1961). To address this question, numerous ecological and evolutionary hypotheses have been developed, with some emphasizing the ability of pathogens and enemies to create and sustain diversity within host or exploited populations (Ehrlich and Raven, 1964; Petermann et al., 2008; Benítez et al., 2013). While pathogens may generate and maintain host diversity, there remains a need to understand the underlying forces and how the processes of creation and maintenance of diversity are interconnected. Therefore, identifying the specific and particular conditions under which pathogens create and maintain hosts diversity may help to disentangle the link between the creation and the maintenance of diversity, from the ecological to the evolutionary perspectives.

Pathogens as key drivers in the emergence of host diversity, particularly intraspecific diversity or polymorphism, can involve various evolutionary processes. Indeed, while hosts and pathogens may co-evolve in an arms race, sometimes leading to diversification in one or in the other species (Yoder and Nuismer, 2010; Best et al., 2009), hosts may also diverge under the selective pressure imposed by non-evolving pathogens. Boot, Milers and collaborators in their theoretical works reported that, under a fecundity-immunity trade-off, a non-evolving pathogen may generate polymorphism in its host population (Boots and Bowers, 1999; Miller et al., 2005; Boots and Haraguchi, 1999). This was possible because a resistant host strain may control and reduce pathogen proliferation in the population, which allows a susceptible host strain to emerge. Empirically, it has been evidenced in different populations of *Pseudomonas* bacteria that, from a single bacteria phenotype, multiple other phenotypes may emerge in the presence of a bacteriophage (a bacteria virus), serving as a selection pressure in the population. In these cases, not only a resistant morph emerged next to the sensitive one, as a consequence of fitness trade-off between exploiter resistance and competitive ability (Brockhurst et al., 2004), but also polymorphism in resistance to the phage is observed, with each resistant morph carrying the cost of its corresponding resistance level (Brockhurst et al., 2005). Other examples of natural enemies (pathogens and predators) creating diversity in their exploited populations have been observed in the phenotypic plasticity in the cotton aphid (Mondor et al., 2005) or in the color polymorphism of other insects such as *Philaenus spumarius* (Harper and Whittaker, 1976) and *Timema* stick (Nosil and Crespi, 2006).

Once polymorphism has emerged, pathogens can play a role in maintaining it through various processes, one of which is the balancing selection they are able to induce within the host population. Balancing selection or balancing polymorphism occurs when polymorphic diversity is preserved due to selective pressures that favor different morphs under varying conditions (Dobzhansky, 1961; Key et al., 2014). Pathogens can drive this process by exerting selective pressures that favor different host morphs over time or in different environmental contexts. One well-described example of this mechanism is found in the Malaria epidemiology in sub-Saharan human populations. Because in the life cycle of the Malaria parasite, human red blood cells (RBC) are exploited by the parasite to replicate in the host, humans with sickle-like RBC are found to better resist to pathogen entry compared to humans with normal cells. Despite the severe anemia caused by sickle-like RBC in hosts carrying the corresponding genotype, the genotype could still be maintained due to the protective effect it offers against the parasite (Elguero et al., 2015; Williams et al., 2005). In some documented cases, the long-term maintenance of polymorphism in the population may further drive the diverse morphs to diverge into different species (Gray and McKinnon, 2007; Hugall and Stuart-Fox, 2012). Thus, the maintenance of polymorphism or intraspecific diversity could serve as the first step to diversification or speciation.

If, as suggested above, pathogens and natural enemies play a central role in fostering intraspecific polymorphisms and promoting interspecific diversity, they may also actively contribute to the maintenance of this interspecific diversity. Indeed, in forest ecosys-tems, Connell and Janzen propose that natural enemies, such as specialist herbivores and pathogens, tend to congregate near their host trees to exploit their seeds and resources (Janzen, 1970; Connell, 1971). As a result, seeds are more likely to survive and germinate farther away from the parent tree, where their natural enemies are less prevalent. This leads to the recruitment of different species around the parent tree (spatial turnover), since the enemies that thrive near the host are not well-adapted to exploit these newly recruited heterospecies. The Janzen-Connell hypothesis has been evidenced at different scales, from the individual levels with fungal and soil-borne pathogens infecting the tree species *Pleradenophora longicuspis* (Bagchi et al., 2010), *Castanopsis fissa* (Liu et al., 2012a), *Prunus serotina* (Packer and Clay, 2000) and *Ormosia glaberrima* (Liu et al., 2012b) to the community (Klironomos, 2002; Hülsmann et al., 2024) and even macroecological scales (LaManna et al., 2017).

Interspecific diversity may also be maintained when different species face a trade-off between competing for resources and defending themselves against natural enemies. Indeed, similar to the mechanism exposed previously in the emergence of polymorphism, the “Killing the Winner” hypothesis suggests that in a resource-limited environment, competition versus defense abilities (against predators or pathogens) are in trade-off such that competitive species may faster proliferate and replace defensive species in a predator- or pathogen-free environment. However, if present, predators, and pathogens may stabilize the system by reducing the density of the dominant species, thus promoting species coexistence (Brockhurst et al., 2006; Winter et al., 2010). This has been highlighted and developed by some theoretical works who described how in an apparent (indirect) competition, shared natural enemies may help to down regulate the more dominant or more abundant species, allowing species to coexist (Holt, 1977; Holt and Lawton, 1994; Holt and Polis2’, 1997; Holt and Bonsall, 2017; Leibold, 1996). In their empirical review on the maintenance of plant diversity by pathogens, Bever et al. (2015) reviewed how pathogens with density-dependent transmission can promote the coexistence of plant species when the competitively superior plant is more susceptible to the pathogen. In multi-plant and multi-pathogen systems, they highlighted that pathogen specialization on specific host plants can regulate host populations, allowing the coexistence of diverse plant species.

The aim of this study is to examine the effects of a trade-off between fecundity and a specific immunity component, pathogen transmission, on diversification processes occurring within the host population. Specifically, we will investigate how this fecundity-transmission trade-off influences the emergence and maintenance of host polymorphism, building on previous works (Boots and Bowers, 1999; Boots and Haraguchi, 1999; Miller et al., 2005). Beyond this, we will explore the connection between the initial emergence and long-term maintenance of polymorphism, the potential for subsequent diversification events, and the maximum number of phenotypes that can be sustained within the system. We will use an eco-evolutionary model developed in our previous study (Djahoui et al., 2025), which incorporates a resource allocation axis into the evolutionary trade-off under consideration. In this framework, only the host population is assumed to evolve. We will base our analysis on the adaptive dynamics approach, while systematically manipulating the strength of the trade-off. The objective is to determine whether such a model construction reveals new or previously unrecognized mechanisms driving pathogen-induced host diversification, with a particular emphasis on intraspecific diversity. We hypothesize that host diversification, if it occurs, will arise under intermediate trade-off strengths, as these are likely to provide an optimal balance between the fecundity and transmission functions. We observe that host diversification does indeed arise under intermediate trade-offs, but only within highly restricted conditions. Moreover, the resulting diversity is consistently limited, with no more than two coexisting host morphs.

## 2 Material and methods

### 2.1 The epidemic model

We use an epidemic model in which the host population is structured into 3 compartments (susceptible, infected and recovered), as shown on figure 1, and given by the following set of differential equations:

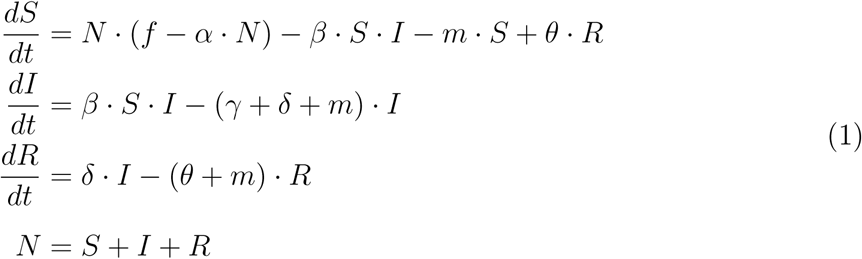

where *S*, *I*, and *R* denote the susceptible, inflected, and recovered densities respectively, and *N* is the total density of the host population. Disease propagation is horizontal, so that newborns are all susceptible and produced according to the classical logistic equation. In this logistic equation, *f* stands for the *per capita* host fecundity rate, *m* the natural host mortality rate, and *α* the host *per capita* intraspecific competition rate. Infection dynamics are captured by the other parameters of the model: *β* the pathogen transmission rate, *γ* the pathogen-induced host mortality rate or pathogen virulence, *δ* the host recovery rate and *θ* the rate of immunity loss in the host population. Analyses of the ecological dynamics can be found in (Djahoui et al., 2025), where the ecological equilibrium states as well as the conditions governing transitions between these equilibrium states are described.

**Figure 1:**
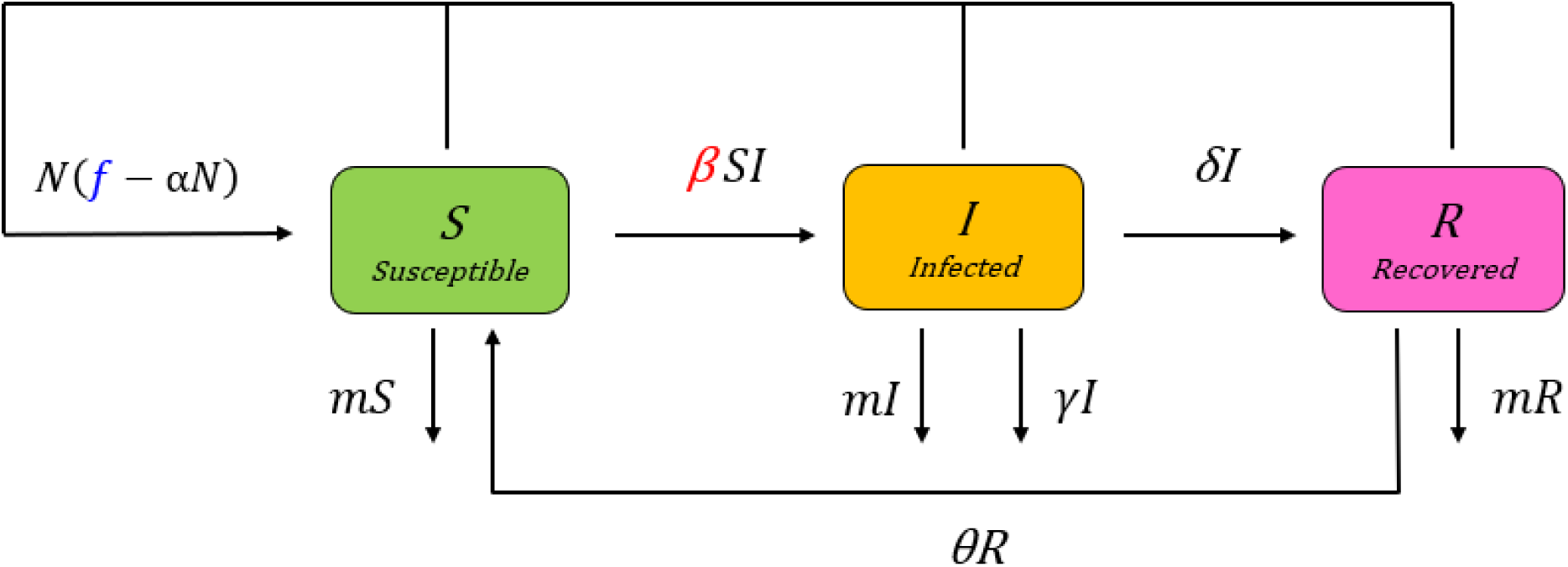
This figure shows the epidemic model (*S* = susceptible, *I* = infected, and *R* = recovered) given by the set of differential equations of equation 1. Arrows indicate the input and output of individuals for each compartment. Pathogen transmission *β*, colored in red, is the immunity component that will be linked to host fecundity *f*, colored in blue, in trade-off formulation.

### 2.2 The fecundity-transmission trade-off

The fecundity-transmission trade-off we use here is based on a resource allocation constraint between host fecundity and immunity, as implemented in (Djahoui et al., 2025). We assume that reducing pathogen transmission in an immune host comes at the cost of its fecundity. This fecundity-transmission trade-off relationship is here modeled along a resource allocation axis, denoted as *x*, representing a continuum of possible allocation strategies. The value of the trait *x* along this axis influences the balance between maximizing reproductive output (when *x* is close to *x_f_*) and enhancing immune function (when *x* is close to *x_β_*), as illustrated on Figure2.A.

Fecundity *f* is given by the following upward Gaussian-shape curve:

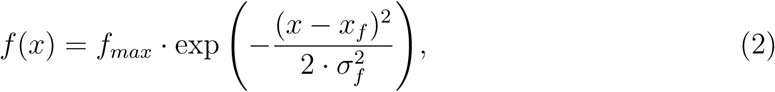

where *f_max_* represents maximum fecundity, *x_f_* the phenotype allowing this maximum fecundity and *σ_f_* a parameter that affects the overlap of the fecundity and immunity functions, translating how fast reproduction is impacted when phenotype *x* evolves toward higher immunity.

Transmission *β* is similarly modeled using a downward Gaussian function:

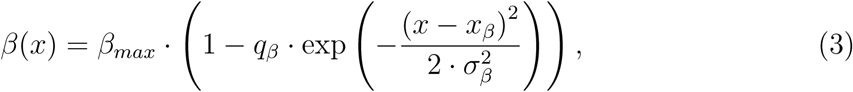

where *β_max_* is the maximal transmission rate happening when host immunity is at its weakest, i.e., far from *x_β_*. At position *x_β_*, the host phenotype invests maximally in immunity, so that pathogen transmission is minimal. The fraction *q_β_* represents the immune system’s effectiveness in reducing transmission at this optimal immunity position, while *σ_β_* constrains how much departing from high immunity, phenotypes will lead to higher transmissions (high *σ_β_* leading to lesser effects).

The strength of the fecundity-transmission trade-off depends on two aspects. First, the distance between the points of optimum fecundity and optimum immunity, represented by *|x_f_ − x_β_|*, which determines the interval on which the trade-off effectively acts. This distance can be adjusted by changing the position of the optimum immunity *x_β_*, while keeping the position *x_f_* for optimum fecundity at 0. Second, the relative spread or overlap of the two Gaussian functions, as this constrains how much a departure from optimal phenotypes, ultimately impact fecundity and transmission respectively. We here fix the spread of the fecundity function, *σ_f_*, to 1 and systematically manipulate *σ_β_*. Note that variations in the two aspects systematically lead to stronger (more concave) or weaker (more convex) trade-offs, as illustrated on figure 2.B, C and D, while the transition areas between concave and convex trade-offs can be considered as intermediate trade-off areas. We will integrate these two aspects into the model analysis to assess whether and how their interaction influences host diversification processes within the system. This approach may uncover mechanisms that drive and maintain diversity within the host population.

**Figure 2:**
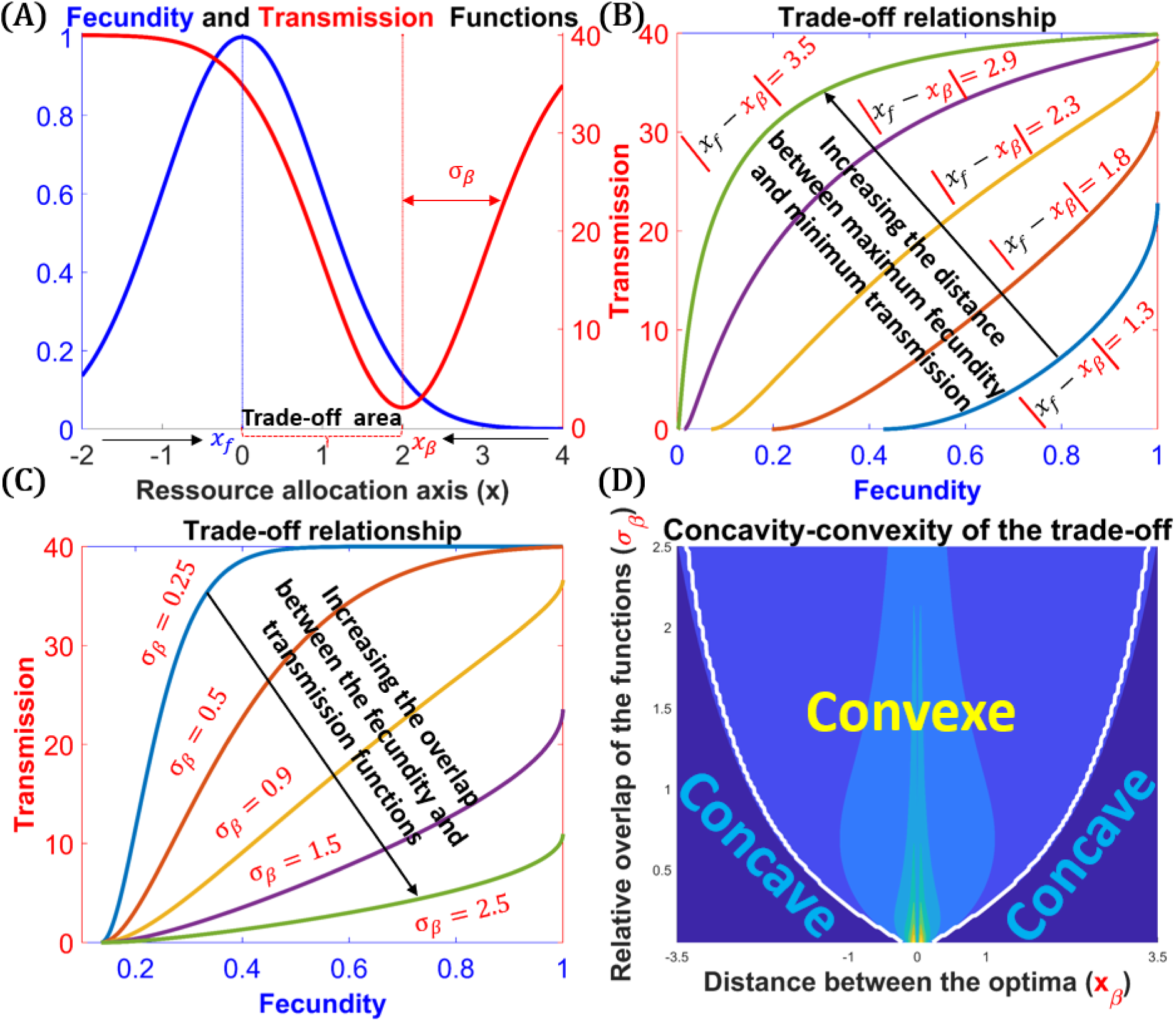
Fecundity-transmission trade-off. **Panel A** shows the fecundity (black curve) and transmission (red curve) as function of the phenotype *x*. The blue and red vertical lines respectively indicate positions of the phenotypes that maximize fecundity and immunity. The interval between these two optima corresponds to the trade-off section, as any variation of *x* in this interval either leads to higher reproduction and lower immunity, or the reverse. Values of *x* outside this interval result in no compromise between fecundity and immunity, leading to directional selection, and the black arrows indicate the direction of evolution starting from these external regions. The host’s position on the trade-off area, along with the overlap between fecundity and transmission (i.e., the overlap of these biological functions), jointly determine the strength of the trade-off. **Panel B** shows the trade-off relationship for various distances between optimum fecundity and optimum immunity (minimum transmission). Increased distances, i.e., increased *x_β_*, lead to concave relationships, thus increasing the trade-off strength. Conversely, decreased distances lead to convex relationships, thus decreasing the trade-off strengths. **Panel C** shows the trade-off relationship for various overlaps between the fecundity and transmission functions. Increased overlap, i.e., increased *σ_β_*, leads to convex relationships, thus reducing the trade-off strength. Again, decreased overlaps increase the trade-off strengths. **Panel D** shows how both the distance between optimum fecundity and optimum immunity and the overlap of the biological functions jointly affect the concavity and the convexity of the trade-off relationship, determined by the mean second derivative along each trade-off relationship. Concave relationships, characterized by negative second derivatives and indicative of strong trade-offs, primarily arise when *x_β_* is significantly increased while *σ_β_* is reduced. Conversely, convex relationships, reflecting weak trade-offs, emerge under conditions of small *x_β_* and large *σ_β_*. Parameters: *β_max_* = 40; *q_β_* = 0.95; *fn* = 1; *x_f_* = 0; *σ_f_* = 01; *x_β_* = 02 and *σ_β_* = 01 for figure A.

### 2.3 Measures of diversification

To explore the conditions underlying host diversification under a fecundity-transmission trade-off imposed by a non-evolving pathogen, adaptive dynamics (AD) emerges as a particularly useful framework (Metz et al., 1992; Geritz et al., 1998; Dieckmann and Ferrière, 2004; Åke Brännström et al., 2013). AD is not only highly effective in modeling eco-evolutionary dynamics (as detailed below), but it also offers the distinct advantage of readily capturing the emergence of polymorphism through evolutionary branching. Our investigation of host diversification therefore directly relates to branching criteria (as detailed below). We will evaluate these diversification criteria by addressing three key aspects: (1) The circumstances under which evolutionary branching can occur; (2) The likelihood of achieving evolutionary branching when multiple attractors exist, determined by the basin of attraction; and (3) The intensity of disruptive selection following the branching events.

Under specific assumptions, including a clonal reproduction and a large size of the evolving population, as well as the rareness, randomness and a very small phenotypic effect of the mutations (Metz, 2012), AD provides a robust mathematical tool for analyzing host evolutionary dynamics, with a particular focus on host diversification. In AD, a rare mutant host with trait value *x_m_* may invade and replace the resident host population with trait value *x* and at its demographic equilibrium, if the mutant invasion fitness *ω*(*x_m_, x*) is positive (Geritz et al., 1997). As in our structured model, each compartment may contribute to the mutant invasiveness, we will use a Jacobian matrix method to account for all these contributions and derive the *ω*(*x_m_, x*). From the Jacobian matrix of the mutant system, this method identifies the largest eigenvalue of the system as a proxy (sign equivalent) of the mutant’s invasion fitness. The full mathematical derivation of *ω*(*x_m_, x*) can be found in (Djahoui et al., 2025).

Successive invasions and replacements of the resident host by the mutant host lead host evolution towards or away from its evolutionary singularities, while the canonical equation given by:

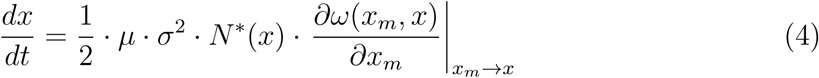

provides both the speed and the direction of such host evolution. In equation 4, *µ* is the mutation rate, *σ*^2^ is the variance brought by the mutational process and *N* ^∗^(*x*) is the density of the resident host population with trait *x* and at the ecological equilibrium.

While the term *∂ω*(*x_m_, x*)*/∂x_m_*, known as the selective gradient of host evolution, is the term effectively influencing the direction of evolutionary changes in the host traits, its nullification identifies the singular strategies in host evolution (Dieckmann and Law, 1996).

Two important criteria are used to characterize a given singular strategy: the convergence criterion and the stability criterion (Eshel, 1983; Eshel et al., 1997; Christiansen, 1991). The convergence criterion evaluates the attractiveness of the singular strategy to nearby phenotypes (convergent if attractive, non-convergent if repulsive), while the stability criterion assesses the invasiveness of the singularity by mutants close to it (invasible or non-invasive). Based on these two criteria, four different types of singular strategies are defined: a continuously stable strategy (CSS), which is convergent and non-invasible, and represents the end point of evolution; a Repellor, which is non-convergent and invasible; a Garden-of-Eden, which is non-convergent and non-invasible; and a branching point, which is convergent and invasible.

Evolutionary branching points are evolutionary attractive singularities on which the evolving host undergoes a disruptive selection. Typically, any nearby mutant close to this point will evolve in the direction of the point where the mutant and the resident hosts may mutually invade and eventually coexist without mutual exclusion, leading to the emergence of polymorphism (Metz et al., 1992; Geritz et al., 1998; Kisdi, 1999).

Since an evolutionary branching point may not be the only one evolutionary convergent strategy in host evolutionary dynamics, host evolution, depending on its starting point, may converge to different evolutionary strategy points, as shown on figure 3.A. For this reason, when multiple evolutionary attractors exist, it is useful to define the probability that host evolution converges to its branching point, which provides the likelihood that the host will undergo a disruptive selection during the course of its evolution. This probability *P* is calculated by dividing the basin of attraction (the set /length of trait values around the branching point that may lead host evolution to converge to that point) by the trade-off area (the region between the optimum positions for fecundity and immunity, within which host evolution is studied). *P* is then given by:

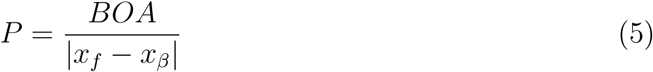

where *BOA* stands for ”Basin Of Attraction”.

**Figure 3:**
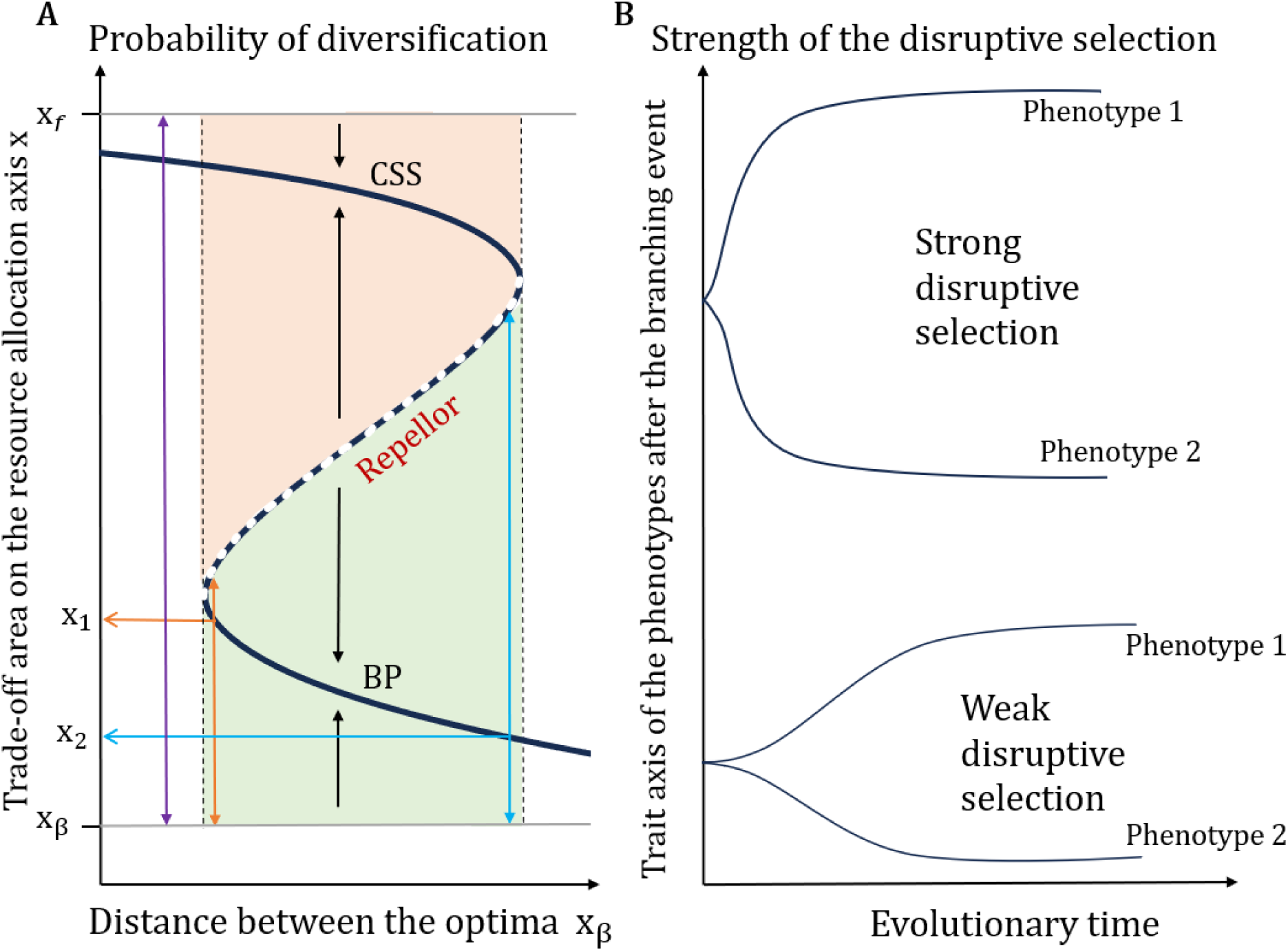
Probability of diversification and strength of the disruptive selection. **Panel A** depicts the basins of attraction of two evolutionary convergent strategies -a CSS and a Branching Point (BP)-separated by an evolutionary non-convergent strategy, here a Repellor. Phenotypes *x_f_* and *x_β_* allow maximal fecundity and immunity, respectively, and they delimit the trade-off area (illustrated by the two-headed purple vertical arrow). The black dashed vertical lines delimit the trade-off strength (here only under various *x_β_*) for which alternative convergent strategies are possible. The black vertical arrows indicate the direction of evolution, while the two-headed orange and light-blue vertical arrows represent the basins of attraction of two branching points for two different trade-off strength (*x_β_*). The orange and light-blue horizontal arrows indicate the trait *x* of the corresponding branching point. Branching probability is calculated by taking the basin of attraction of the branching point that we divide by the size of the trade-off area. **Panel B** illustrates branching processes under two different strengths of disruptive selection. With strong disruptive selection, two distinct host morphs with widely spaced trait values emerge over a relatively short evolutionary timescale. Conversely, weak disruptive selection results in the emergence of host morphs with closely aligned trait values, occurring over a longer evolutionary period.

As an illustrative example, figure3.A shows the probability of diversification of two distinct branching points occurring at different distances between the optima in fecundity and transmission. Strategy *x*_1_ (low distance) has a lower probability of diversification compared to strategy *x*_2_, as evidenced by its smaller basin of attraction (shown by the two-headed orange vertical arrow).

At a given branching point, the mutual invasibility of the mutant and resident hosts characterizes the point as an evolutionary instable point, where the condition of this instability is given by:

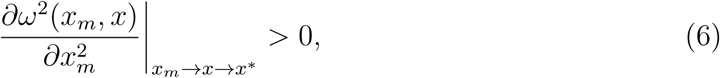

with *x*^∗^ being the trait value at the singularity.

Equation 6 is actually the second derivative of the mutant invasion fitness at the exact branching point with respect to the mutant trait value (Åke Brännström et al., 2010). It quantifies the intensity of disruptive selection, representing the strength of the selection pressure driving the host population to diverge into two distinct morphs. Higher values of Equation 6 imply strong disruptive selections, while lower values imply weak disruptive selections, as shown on figure3.B.

Figure 4.A depicts in details an example of the host evolutionary dynamics under a particular trade-off strength generating three evolutionary strategies, a CSS, a Repellor and a branching point, corresponding respectively to the black, the red and the blue dots on figure4.A, along with the direction of evolution (one-headed arrows) and with the basins of attraction (BOA) of the attractive singularities (two-headed black arrow for the CSS and two-headed blue arrow for the branching point). This figure is a classical pairwise invasibility plot (PIP) on which mutant fitness is represented, where positive regions (colored in green) are the regions where a mutant host having a positive invasion fitness may successfully invade and replace the resident host population. As the BOA of the branching point is larger than that of the CSS, there is a higher probability for the host to evolve to the branching point than to the CSS, at least under this particular set of parameters and conditions. Since the evolutionary strategy that will be reached will depend on the starting point of evolution, figure4.B shows how, starting from two different points, the host evolution converges either to the CSS or to the branching point where the host diverges into two separate morphs. This figure is a simulation of host evolutionary dynamics under stochastic mutations. When only the mutant invasion fitness is positive, the mutant replaces the resident while when the mutant and resident invasion fitness are both positive, host dimorphism is allowed. As here the CSS is very close to the maximum fecundity, we suggest that, in a fecundity-transmission trade-off, if at the starting point of evolution, the host bears a high cost of pathogen transmission without enough fecundity, then it evolves to the maximum fecundity despite the pathogen harm whereas when the investment in fecundity and in immunity is quite balanced, the host may evolve to the branching point.

**Figure 4:**
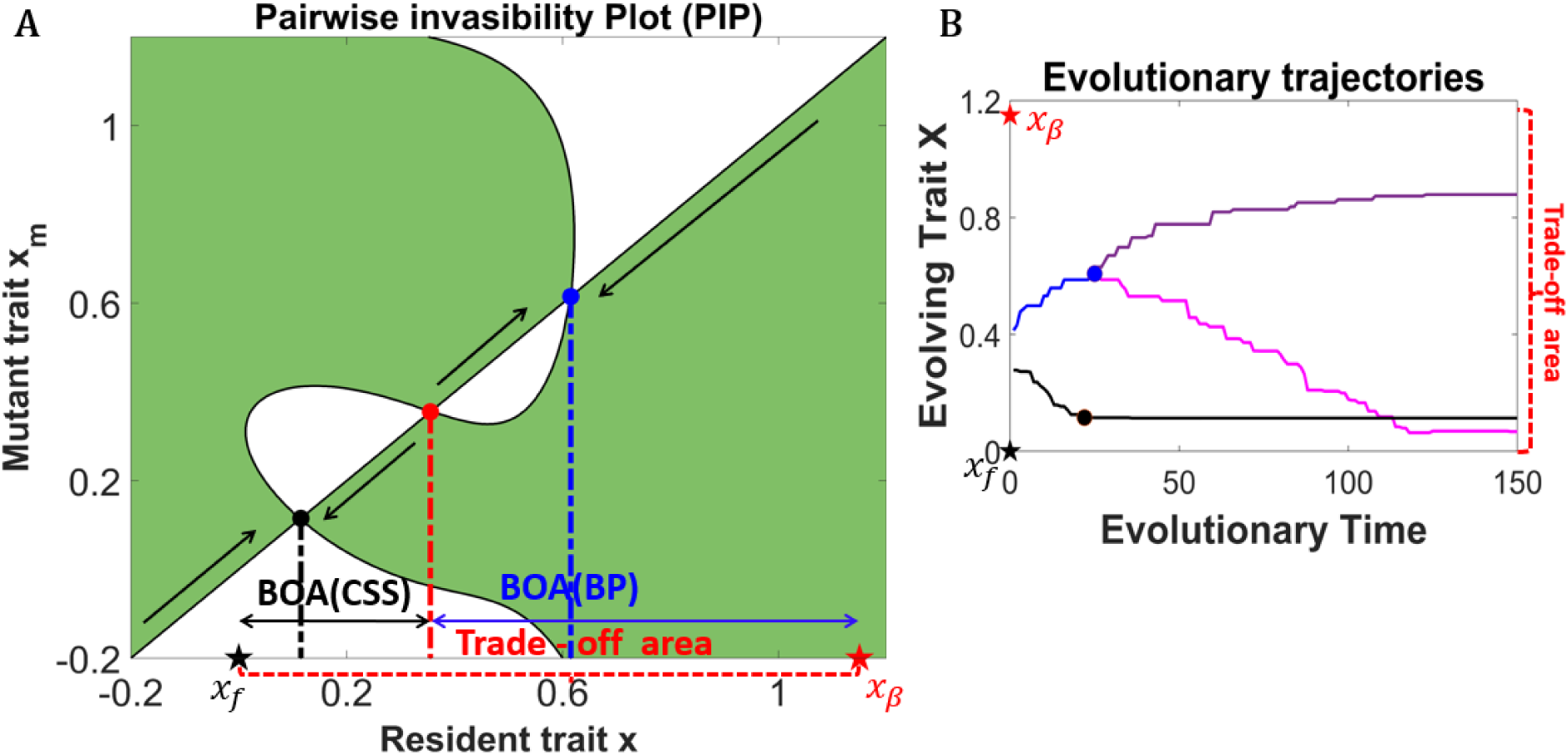
**Panel A** is a pairwise invasibility plot (PIP) showing the evolutionary outcome of a mutant host successfully invading and replacing the resident host population through small mutational steps. While mutants can invade and replace residents in the colored region where the mutant fitness is positive, successive invasions and replacements lead host evolution. This PIP shows multiple alternative and convergent strategies (as in the yellow region of fig 5.A: a CSS (black dot) and a branching point (blue dot), separated by a Repellor (red dot). The one-headed black arrows indicate the direction of evolution. **Panel B** depicts the evolutionary trajectories of host evolution and shows that, depending on the starting point of this evolution, the evolution may converge either to the CSS or to the branching point. The black curve in the trajectory of the host evolving to the CSS, whereas the blue curve is the trajectory of the host evolving to the branching point. The lilac and purple curves are the trajectories of the two host phenotypes after the branching event. Parameters: *x_β_* = 1.15, *σ_β_* = 0.4, *β_max_* = 40, *r_β_* = 0.95, *x_f_* = 0, *σ_f_* = 1, *f_max_* = 1, *γ* = 10, *δ* = 1, *θ* = 1, *m* = 0.1, *α* = 1.

## 3 Results

### 3.1 Pathogens rarely promote the emergence of host polymorphism

Our analysis indicates that pathogens hardly enable the emergence of diversity. Indeed, diversification in observed only within a narrow range of trade-off values transitioning between concave and convex shapes, corresponding to intermediate trade-off strengths, thus aligning with our prediction. When diversification is possible, its probability of occurrence is low and the associated force of disruptive selection is generally weak. When no diversification occurs, the host evolutionary outcome is always a CSS, while under host polymorphism, a highly fecund and a highly resistant strategy may coexist.

Pathogen-induced host polymorphism occurs in a restricted domain (figure 5.A) and under particular conditions. Since the concavity or convexity of the trade-off relationship defines its strength, polymorphism tends to emerge near the transition zone between concave and convex trade-offs. This transition zone likely represents an optimal balance between investments in fecundity and immunity, where reduced pathogen transmission is achieved without excessive reproductive cost.

**Figure 5:**
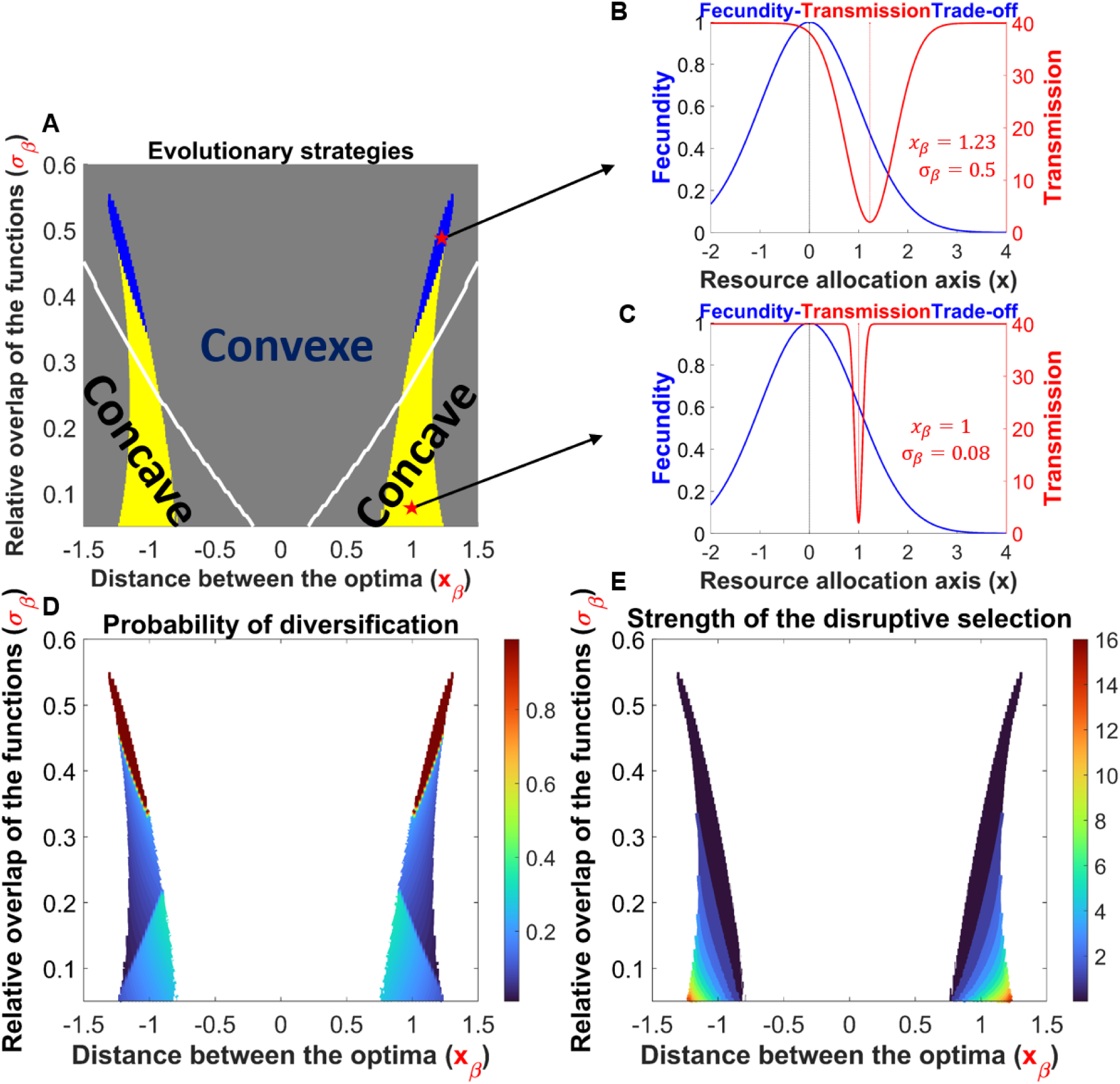
**Panel A** depicts the evolutionary strategies for various trade-off strengths, represented by both the different distances between the positions for optimal fecundity and optimal immunity (*x_β_*) and the different levels of relative overlap (*σ_β_*) between the fecundity and immunity functions. In the blue area, the host exhibits a single evolutionary strategy, a branching point, while in the yellow area, three evolutionary strategies are obtained (a CSS and a branching point, separated by a Repellor). Gray indicates that no diversification happens, as only CSS are obtained. The white curves indicate the transition between concave and convex fecundity-immunity trade-off relationship. **Panel B and C** show two examples of overlap of the biological functions (moderate and small respectively), and the corresponding host evolutionary outcomes are designated by red stars on panel A. **Panel D** shows the probability of diversification of each evolutionary outcome in the branching area. **Panel E** illustrates the strength of the disruptive selection, which relatively increases under weak overlaps. Parameters: *β_max_* = 40, *r_β_* = 0.95, *x_f_* = 0, *σ_f_* = 1, *f_max_* = 1, *γ* = 10, *δ* = 1, *θ* = 1, *m* = 0.1, *α* = 1. Parameters: *β_max_* = 40, *r_β_* = 0.95, *x_f_* = 0, *σ_f_* = 1, *f_max_* = 1, *γ* = 10, *δ* = 1, *θ* = 1, *m* = 0.1, *α* = 1.

Specifically, regarding the overlaps of the biological functions, one of the parameters investigated as influencing trade-off strength, we observe that low overlaps are crucial for host diversification to arise, as higher overlaps tend to inhibit the emergence of diversity (see figure 5).A. In the regions where the overlaps are sufficiently low to promote polymorphism, we find that for moderate overlaps (relatively larger *σ_β_*, as shown on figure 5.B), this polymorphism emerges in a narrow region of single evolutionary strategy corresponding to a branching point (depicted in blue in Figure 5.A). Conversely, for very low overlaps (relatively smaller *σ_β_*, as shown on figure 5.C), polymorphism emerges across a broader region of parameters, but such parameter sets lead to two distinct, alternative and convergent evolutionary strategies: a continuously stable strategy (CSS) and a branching point, separated by a repulsive strategy, or Repellor (highlighted in yellow on figure 5.A). This illustrates how the various investment strategies toward fecundity and immunity, driven by the differences in the overlaps of their biological functions, may account for the distinct evolutionary outcomes observed. Specifically, very small overlaps translate, for a given host position in the trade-off area, too higher pathogen transmission per infected compared to more moderate overlaps (figure 5.B and C). Therefore, the wider region of host diversification obtained under small overlaps shows how slightly increasing the trade-off constraint affects host evolutionary dynamics by increasing the possibility of polymorphism. Apart from the domain of polymorphism, any other combination of *x_β_* and of *σ_β_* results in a single evolutionary strategy corresponding to a CSS in host evolution. Interestingly, single evolutionary outcomes corresponding to branching points (blue region) typically occur under convex trade-off relationships. In contrast, regions where alternative strategies emerge (yellow region) are predominantly associated with concave trade-off relationships.

When polymorphism emerges, we notice that it is often still unlikely to happen (Figure 5.D). Indeed, the probability of diversification is obviously 1 when only branching points exist (red on Figure 5.D), but when alternative evolutionary strategies exist (Branching and CSS), the probability of diversification is generally low (less than 0.5). Most evolutionary dynamics indeed eventually converge to the CSS. Finally, the speed of the diversification, as measured by the strength of the disruptive selection on figure 5.E, is mainly dependent on the overlap between the fecundity and the transmission functions. Diversification is fast only for very weak overlap of the functions, meaning that high trade-off constraints increase the strength of the disruptive selection, i.e., the branching constraint in the host.

Beyond the shape of the trade-off function, ecological dynamics also affect the possibility of diversification (see Figure 6). However, our results, showing that diversification seldom occurs and primarily around the transition zone between concave and convex trade-off scenarios, remain largely valid. While, for the investigated ecological components, we still obtain two separated regions of polymorphism, we notice that increasing the intraspecific competition *α* reduces the distance between the two regions, while increasing host mortality *m* reduces the length of these regions of polymorphism. Therefore, varying these two ecological components may allow the emergence of polymorphism under relatively different concavities or convexities of the trade-off relationship. Interestingly, high competition combined with elevated host mortality rates make the emergence of host polymorphism challenging.

**Figure 6:**
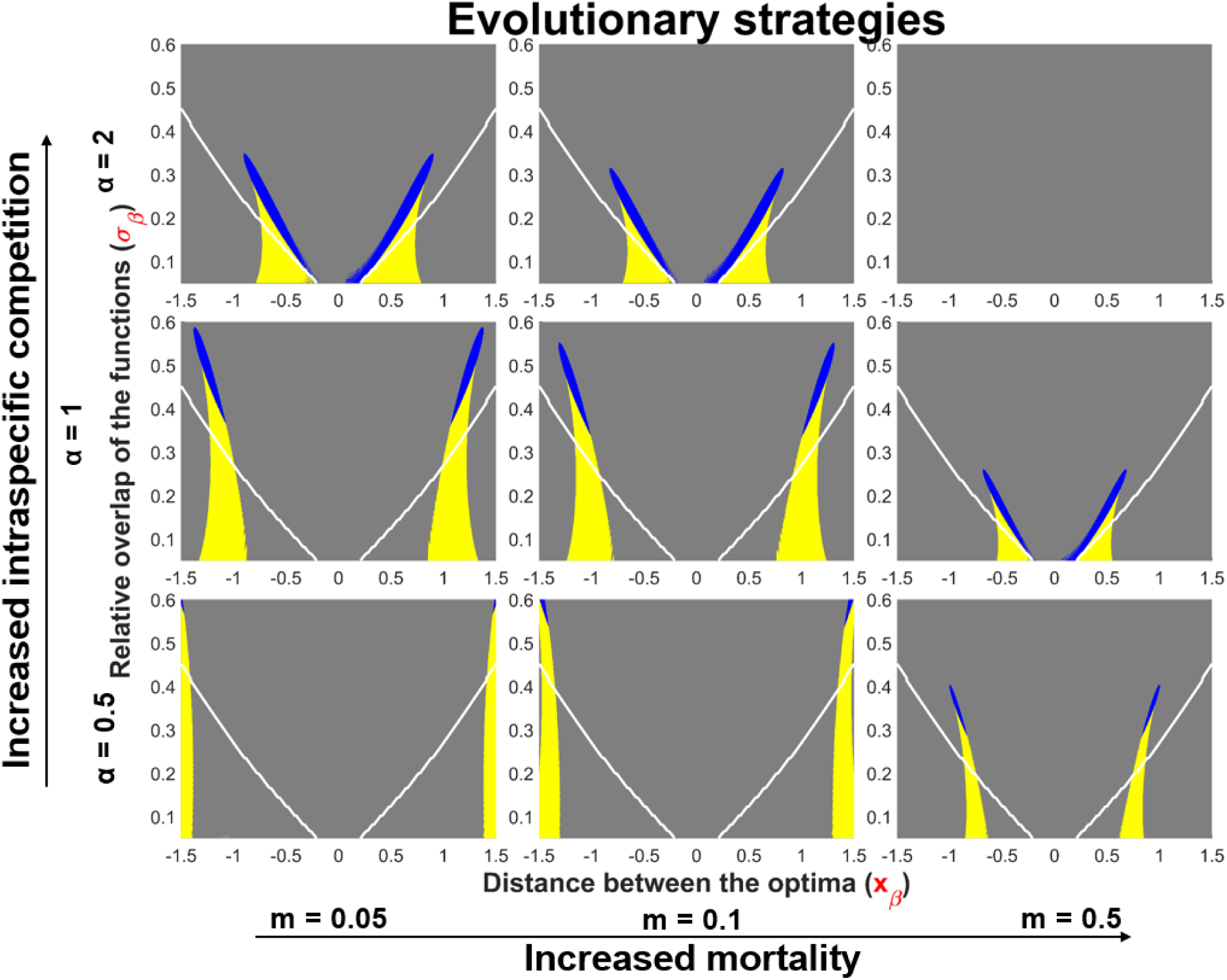
This figure shows the emergence of polymorphism under various intraspecific competition *α* and various host mortality rates *m*. Parameters: *β_max_* = 40, *r_β_* = 0.95, *x_f_* = 0, *σ_f_* = 1, *f_max_* = 1, *γ* = 10, *δ* = 1, *θ* = 1

### 3.2 Each branching event leads to the maintenance of host polymorphism

We assessed the maintenance of host polymorphism after each branching event in the branching area and found that each branching event leads to the maintenance of polymorphism, regardless of the number of singular strategies located in the branching area (one or three) and regardless of the strength of the disruptive selection (see figure S1). Moreover, as exemplified on figure 4.B, polymorphism generally leads to two strategies or morphs, one investing more in fecundity at the expense of immunity and the other prioritizing immunity at the expense of fecundity. Our suggestion is that theses two distinct strategies are maintained because they are approximately equivalent in their respective costs and benefits, otherwise exclusion of one morph by the other may happen. Nevertheless, as this analysis was computational demanding, we could investigate the maintenance of polymorphisms only for parameters displayed on figure 5.A. Since our simulations demonstrate that polymorphism, once emerged, is subsequently maintained, we conclude that, under the biological parameters considered in our analysis, the conditions that facilitate the emergence of host polymorphism are indeed the same as those that support its maintenance.

### 3.3 Subsequent branching events and the maintenance of more than two host morphs are impossible

We now investigated in the possibility of subsequent branching events in the system and found that higher order diversity is impossible. The fact that eco-evolutionary dynamics cannot lead to more than two phenotypes can be readily understood from the ecological system. Indeed, assuming subsequent branching events in the system implies assuming the coexistence of the two host morphs with the new invading mutant from one or the other morph. We therefore question the possibility of subsequent branching events by simply looking at the possibility of the coexistence of more than two host morphs. As-suming that there are more than two host morphs in the system, and that *M*_1_ denotes the system of differential equations of morph 1, which can be written as:

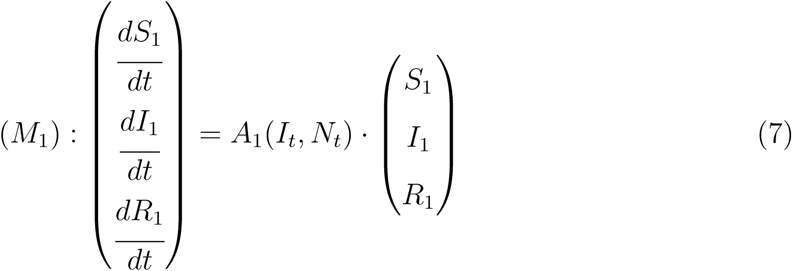

where *A*_1_(*I_t_, N_t_*) is a linear system of the variables *I_t_* and *N_t_*, and given by:

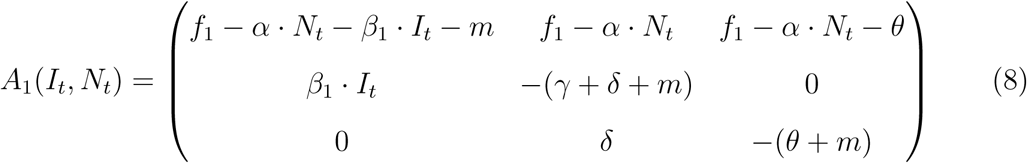

where *N_t_* denotes the total density of all host morphs and *I_t_* is the density of all infected individuals in the system.

Coexistence of at least 3 host morphs is only possible if the following condition holds:

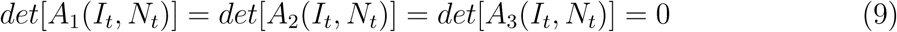

where *A*_2_(*I_t_, N_t_*) and *A*_3_(*I_t_, N_t_*) are the linear systems of variables *I_t_* and *N_t_* for the systems of differential equations of morph 2 and morph 3 respectively. As *N_t_* and *I_t_* are the only two variables present in condition 9, the condition cannot be fulfilled, unless in a degenerate case.

Figure 7 is a graphical illustration of the possibility of the coexistence of two host morphs. Each isocline represents the feasible solution of its corresponding host morph, and the intersection of the two isoclines (red dot) represents the point of coexistence of the two morphs. For three morphs to coexist, the isocline of the potential third morph must intersect the equilibrium point where two morphs coexist. This implies that the coexistence of more than two host morphs is highly unlikely and would occur only under very specific conditions. Consequently, further branching events are not feasible, and the system is expected to support a maximum of two coexisting host morphs.

**Figure 7:**
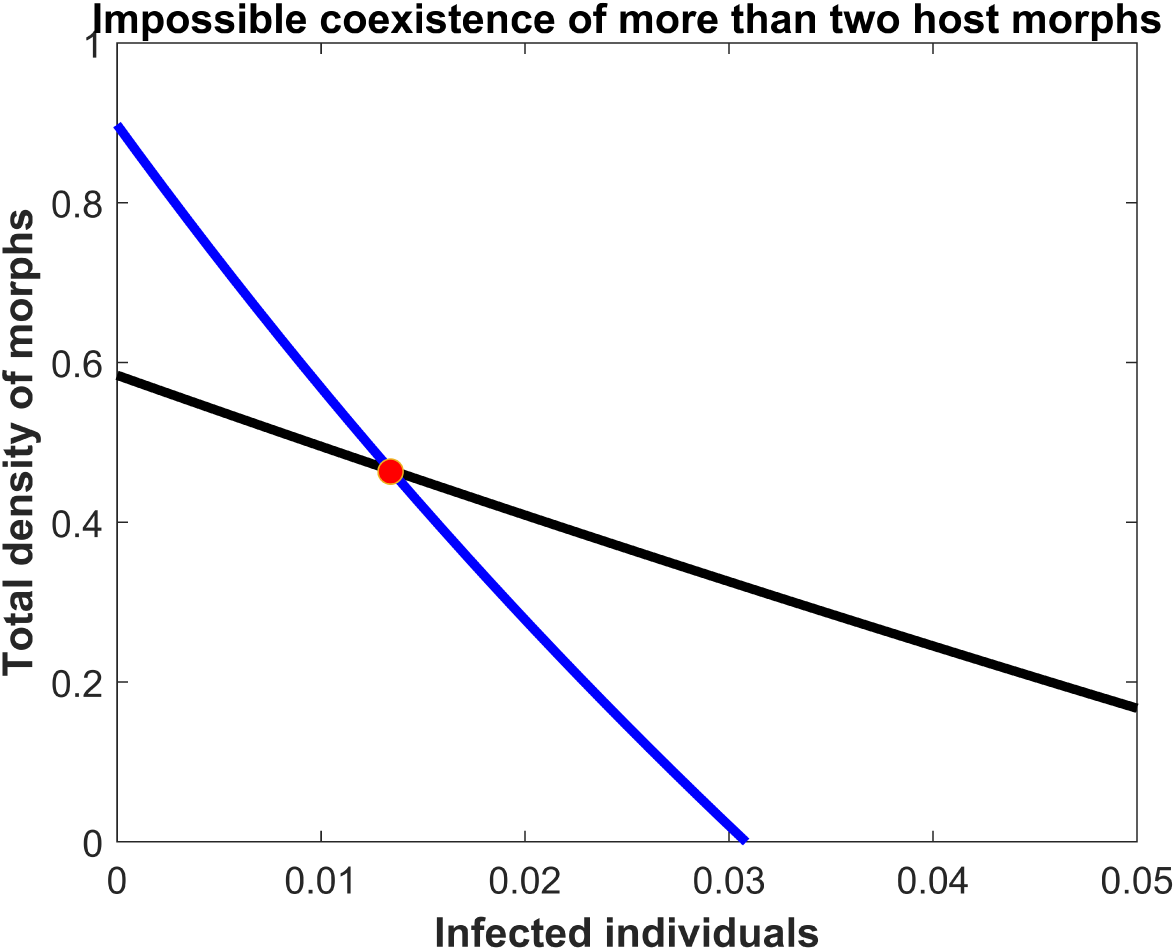
This figure illustrates the ecological feasibility of coexistence between two host morphs. Each isocline graphically represents the conditions necessary for the persistence of its respective morph within the system. The intersection of these two isoclines, marked by the red point, indicates the equilibrium at which the two morphs can coexist. This balance point makes it challenging for a third morph to coexist with the established two, as any additional morph would need to intersect precisely at the two-morphs equilibrium point to be maintained within the system. Parameters: *x_β_* = 1.1475, *σ_β_* = 0.4, *β_max_* = 40, *r_β_* = 0.95, *x_f_* = 0, *σ_f_* = 1, *f_max_* = 1, *γ* = 10, *δ* = 1, *θ* = 1, *m* = 0.1, *α* = 1, *x*_1_ = 0.8714, *x*_2_ = 0.0687.

## 4 Discussion

While pathogens are widely considered an important source of diversification and as allowing the persistence of diversity through various mechanisms, our investigation of host evolution shows that such pathogen-induced polymorphism is quite rare and arises only under specific circumstances. While the distance between optimum positions in fecundity and immunity (*x_β_*) and the overlap of the two biological functions (*σ_β_*) collectively shape the trade-off strength, we notice that only a small region of intermediate trade-off strengths -defined where the fecundity-transmission trade-off switches between concave and convex relationship-are required for this polymorphism to emerge. Interestingly, ecological components such as host intraspecific competition and natural mortality may shift and/or change the size of the domain of polymorphism, thereby allowing polymorphism under relatively weaker or relatively stronger trade-off strengths. While each diversification event leads to the long-term maintenance of this diversity, high diversity is here impossible; diversification always leading to two coexisting host morphs.

Although host polymorphism under pathogen pressure is possible, its rarity and the specific conditions under which it occurs stand in contrast to the common view that pathogens generally promote and sustain host diversity. This discrepancy might have a specific explanation. In our study, the simplified two-species system (one host and one pathogen), while useful for modeling purposes, may not accurately capture the complexity of interactions that typically occur in real-world host-pathogen systems. In natural systems, multiple pathogen species can jointly exert various selective constraints on host populations, potentially amplifying the pressure for diversification. For instance, it has been observed that bacteria and viruses can impose complementary selection constraints on their hosts, such as in the case of the ABO histo-blood group antigen polymorphism in humans, maintained through the combined action of different pathogens (Seymour et al., 2004). Similarly, the genetic polymorphism in toll-like receptor genes involved in innate immunity is preserved in roe deer (*Capreolus capreolus*) due to an antagonistic selection pressure from the intracellular pathogens *Toxoplasma* and *Chlamydia*, reported by (Quéméré et al., 2021). Others reported that multiple parasites are drivers for the class IIB major histocompatibility complex polymorphism in the three-spined sticklebacks (*Gasterosteus aculeatus L.*) (Wegner et al., 2003). While these examples primarily highlight the maintenance of host polymorphism through simultaneous selection pressures imposed by divers pathogens, it is reasonable to speculate that the involvement of multiple pathogens could increase the selection pressure to a point where it not only preserves polymorphism but also enhances the likelihood of the emergence of the polymorphism. This suggests that the simplified model in our study may overlook the more complex ecological interactions that could drive greater diversification in natural systems.

An alternative explanation for the limited pathogen-induced host diversification observed in our study could be tied to the fact that we did not explore the potential antagonistic co-evolutionary dynamics that often occur within host-pathogen systems. These co-evolutionary processes are known to promote or enhance diversification (Yoder and Nuismer, 2010; Hembry et al., 2014; Best et al., 2010), and their absence from our investigation might have influenced the overall findings regarding host diversification. For example, the remarkable diversity of butterfly species and their host plants likely originated from a prolonged process of adaptation and counter-adaptation within this insect-plant system (Ehrlich and Raven, 1964). A similar pattern is observed in bacteria-bacteriophage systems, where coevolution appears to drive the diversity observed in both groups (Koskella and Brockhurst, 2014). Therefore, by not accounting for these interactions, which can play a critical role in driving adaptive changes in both hosts and pathogens, we may have overlooked a key factor that could contribute to greater diversification in host evolutionary dynamics.

Trade-off strength may influence the emergence of host polymorphism. Indeed, while a fecundity (and/or susceptibility)-resistance trade-off, similar to the one investigated here, has been identified as a factor in the emergence of resistant and susceptible strains in a host population with directly transmitted pathogens (Boots and Bowers, 1999) and free-living pathogens (Miller et al., 2005), in the case of directly transmitted pathogens, a high level of resistance combined with either high or low costs of resistance appears to drive the diversification processes observed. Hence, the level of host resistance (or immunity) and the cost of this resistance may constitute important factors determining the emergence of host polymorphism. Our current study, revealing that the transition zone between concave and convex trade-offs, which can here be associated to intermediate trade-offs with balanced investments toward fecundity and immunity, promote host polymorphism, contrasts with the study of Boots and Bowers in which host polymorphism only occurs under high levels of resistance or immunity. One explanation of this divergence could reside in the way we implemented our trade-off, which was not the same in this previous study, and this new approach may affect the measurement of our trade-off strength. This shows the importance of identifying and implementing the specific trade-off mechanisms, with the trade-off characteristic applying to the host-pathogen system of interest, in the analysis of the evolutionary outcome in a given system.

We also find that the conditions for the emergence and for the maintenance of host polymorphism are the same. A simple explanation of this could be that whereas pathogens generate polymorphism in their hosts through the selection pressure they impose on the hosts, the constant selection pressure of pathogens in an environment with polymorphic hosts may help to the maintenance of this polymorphism. While Rainey and collaborators reviewed that in bacteria populations, the conditions of the emergence of polymorphism, including ecological opportunity and competitive trade-offs, also account for the maintenance or the stability of this polymorphism (Paul B. Rainey and Travisano, 2000), Brännström and collaborators showed that in a food-web model, the conditions that favor the emergence of diversity do not necessarily promote the maintenance of the diversity in evolved communities, and *vice versa* (Åke Brännström et al., 2010). Therefore, while we here observe that the emergence and the maintenance of host polymorphism are positively linked, this positive link may be restricted to particular systems and may thus change, depending on the evolving system considered.

Two host morphs may coexist after the first branching event. However, a subsequent branching event is impossible because the coexistence of three morphs is not feasible. Our explanation of this is that the condition of coexistence of three morphs, shown in equation 9, depends on the density of infected individuals and on the total density of all host morphs. Because this condition involves only these two densities, it limits the possibility of introducing a third morph into the system. This can be assimilated to a previous hypothesis suggesting that the number of coexisting species in a system must not exceed the number of available resources (MacArthur and Levins, 1964) or the number of limiting factors (Levin, 1970; Armstrong and McGehee, 1980). Thus, the two densities, mentioned here, can be viewed as limiting factors that may restrict the number of coexisting species in the system. Therefore, a third factor is here required for the coexistence of three morphs to be possible.

While our results show that pathogens may generate and maintain polymorphism in their host populations through a fecundity-transmission trade-off imposed on the host, our study could be deepened with additional investigations. Indeed, in our analysis, we only focus on pathogen transmission as an immunity component in the trade-off formulation, with the idea that immune hosts are able to block pathogen transmission within their populations. In reality, there are numerous trade-off components that can be isolated from our model. Through its immune response, the host may enhance its recovery rate, reduce pathogen virulence, or delay the loss of immunity. However, we limit our analysis to the fecundity-transmission trade-off due to the time-consuming nature of our simulations, which makes it challenging to explore other immune components. Nonetheless, it would be valuable to investigate novel fecundity-immunity trade-offs that involve these additional immune factors and explore their evolutionary implications on host diversification. This could help to identify and classify immunity components that contribute to generating and maintaining host polymorphism from immunity components that do not contribute to polymorphism or have different evolutionary consequences.

Our results show that, under particular conditions, pathogen may induce polymorphism in their host population. Although this polymorphism is here rarely observed, our results enable to disentangle a possible hidden mechanism of this pathogen-induced intraspecific host diversification and the maintenance of this diversity. Recall that in our model, there is no ecological niche differentiation between the resident host and the new invading mutant host. This means for instance that resident and mutant hosts exploit the same resource, on the same habitat or territory and at the same time (Barabás et al., 2018). In this case, host diversification and the maintenance of this diversity should be impossible according to the Gause principle of competitive exclusion (Kneitel, 2008). Therefore, the diversification event observed here during the evolutionary course, as well as the maintenance of this diversity, are certainly induced by the presence of the pathogen, thereby showing its potential ability to generate and/maintain biological instead of ecological niche differentiation between two host phenotypes present on the same area. This biological niche differentiation is reflected in the specialization of the host morphs on one specific biological function, fecundity, or immunity (reduce pathogen transmission), which also allows host morphs to coexist even though morphs share the same ecological niche. Our results thus provide a better understanding of how natural enemies may promote and maintain diversity within their exploited populations.

## Acknowledgments

We would like to thank Dr. Thomas Koffel for his valuable contributions to the discussions of this manuscript.

## Supplementary material

### S0.1 Maintenance of the two-morphs diversity

**Figure S1:**
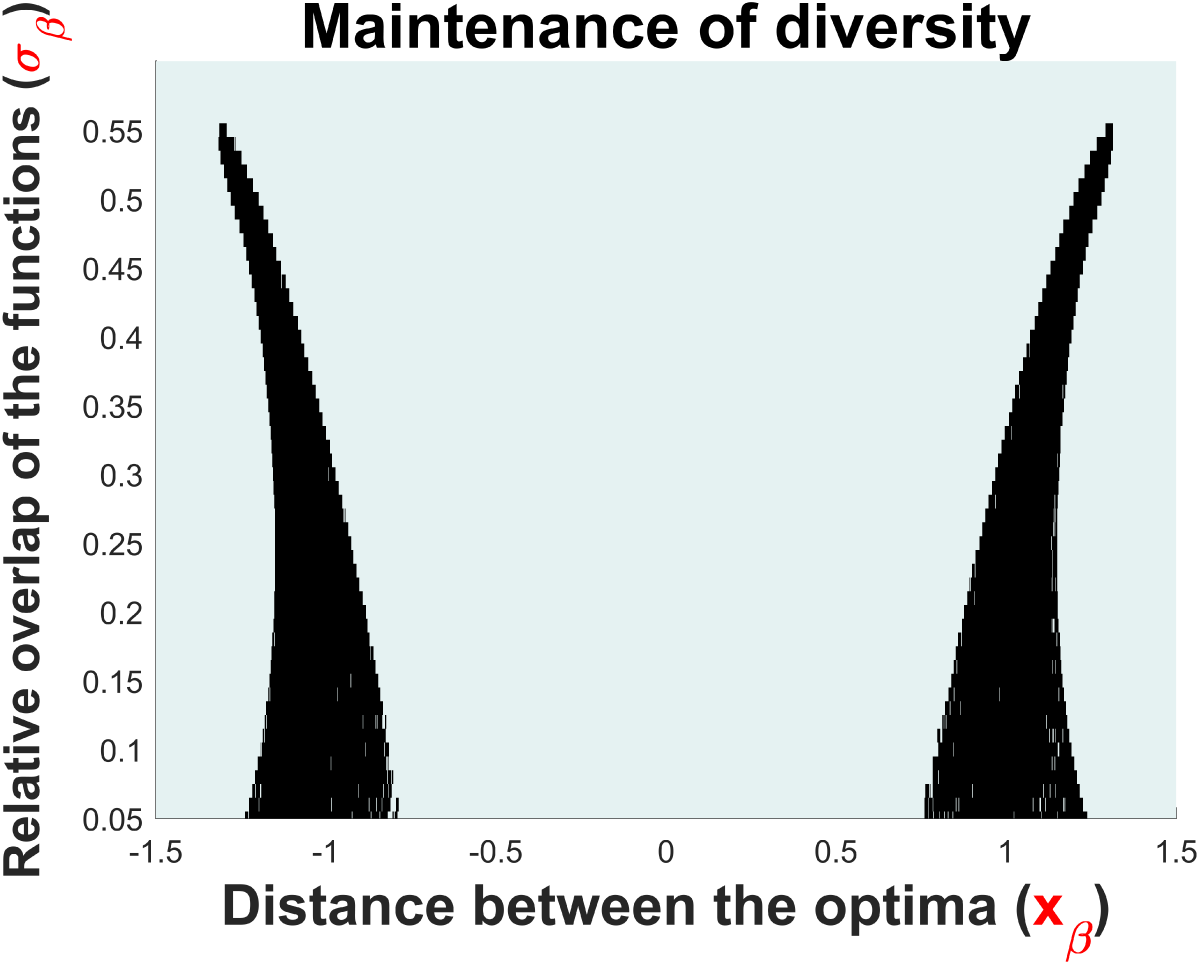
This figure illustrates the long run coexistence of two morphs in the system after each branching event in the branching area. It shows that all branching events lead to the maintenance or the coexistence of the two emerged host morphs. Parameters: *β_max_* = 40, *r_β_* = 0.95, *x_f_* = 0, *σ_f_* = 1, *f_max_* = 1, *γ* = 10, *δ* = 1, *θ* = 1, *m* = 0.1, *α* = 1.

In our investigation in the maintenance of diversity after the first branching event, we followed the densities of both host phenotypes after each branching event and for a sufficient evolutionary time, allowing the system to potentially reach its eco-evolutionary equilibrium. When at the potential eco-evolutionary equilibrium, both morphs remain present, then polymorphism is accepted, while when one or both morphs go extinct (a density below the critical value of 10^−^6), polymorphism is rejected. The simulations were performed for each combination of *x_β_* and *σ_β_* in the branching area, and details are presented in S0.2.

### S0.2 Matlab codes for figure 4.B, figure 5.A, D and E, and figure S1

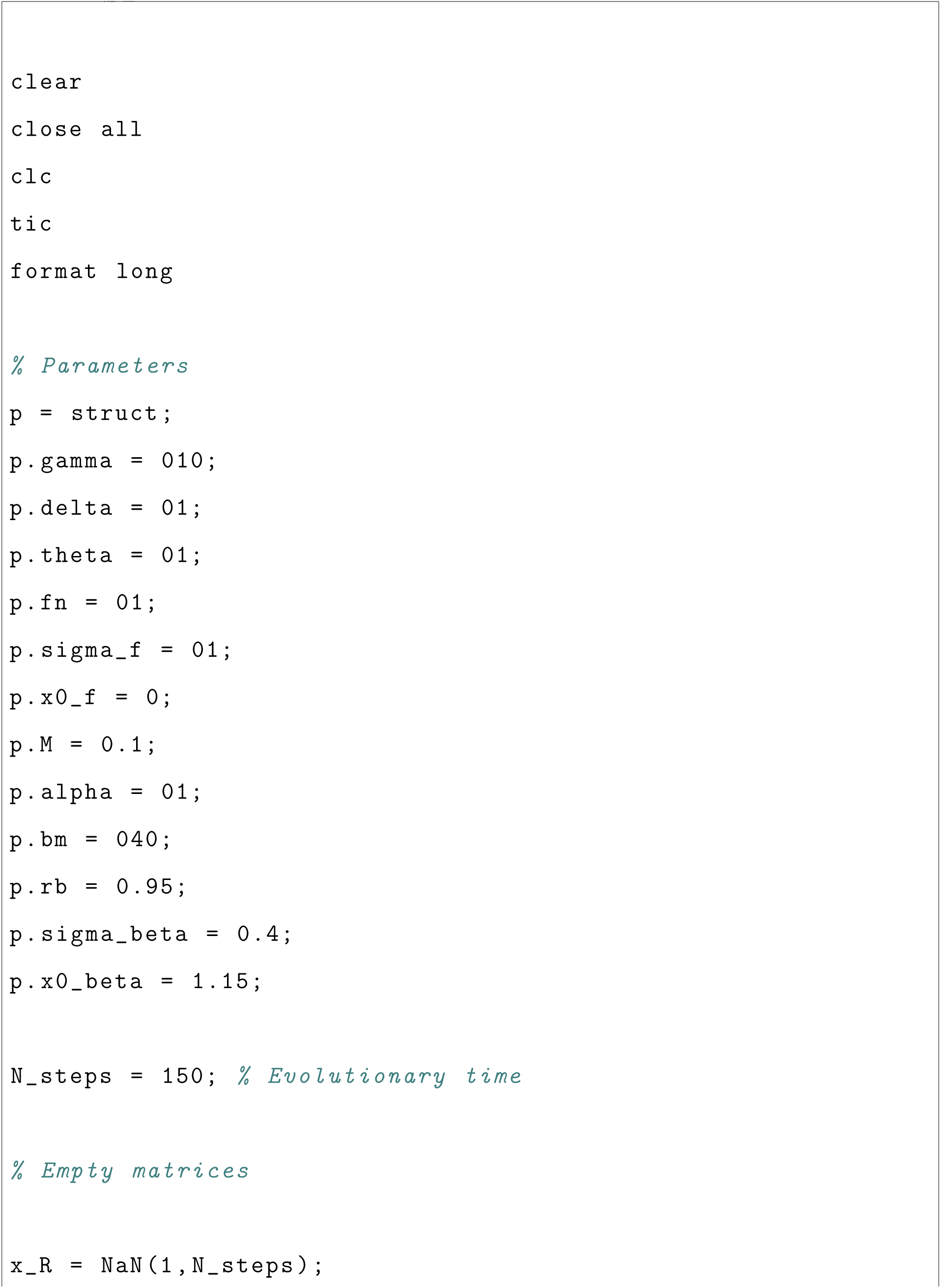
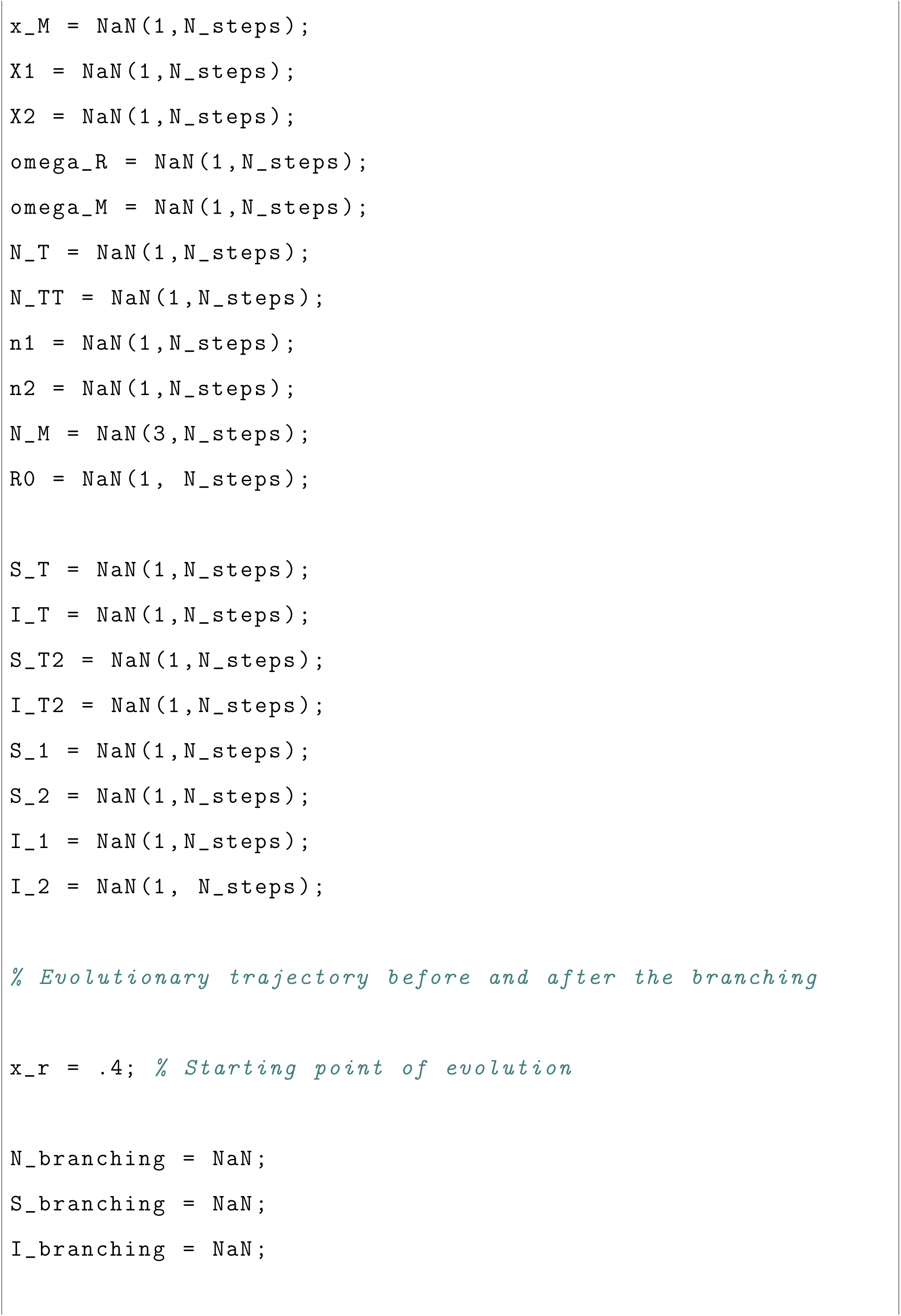
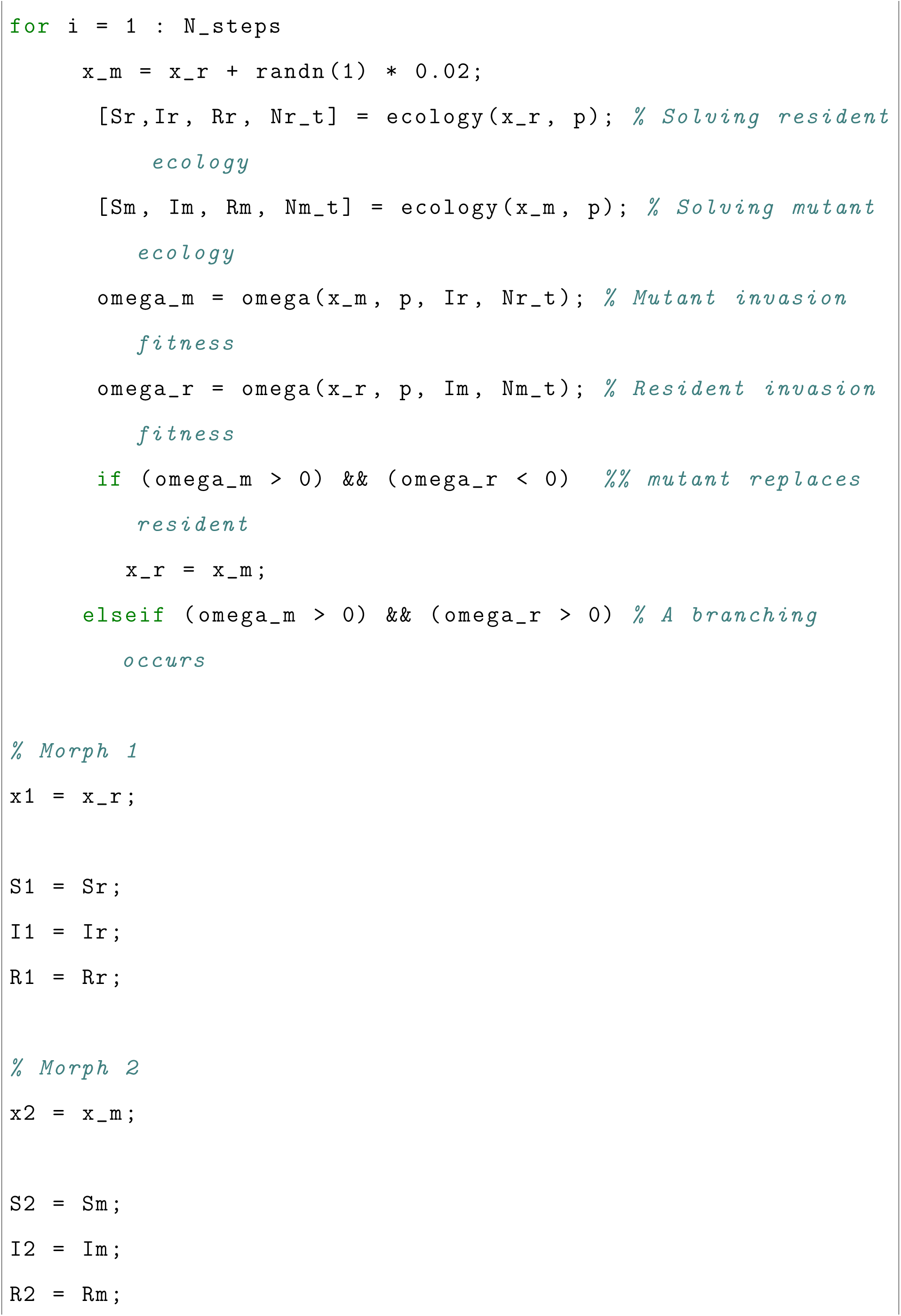
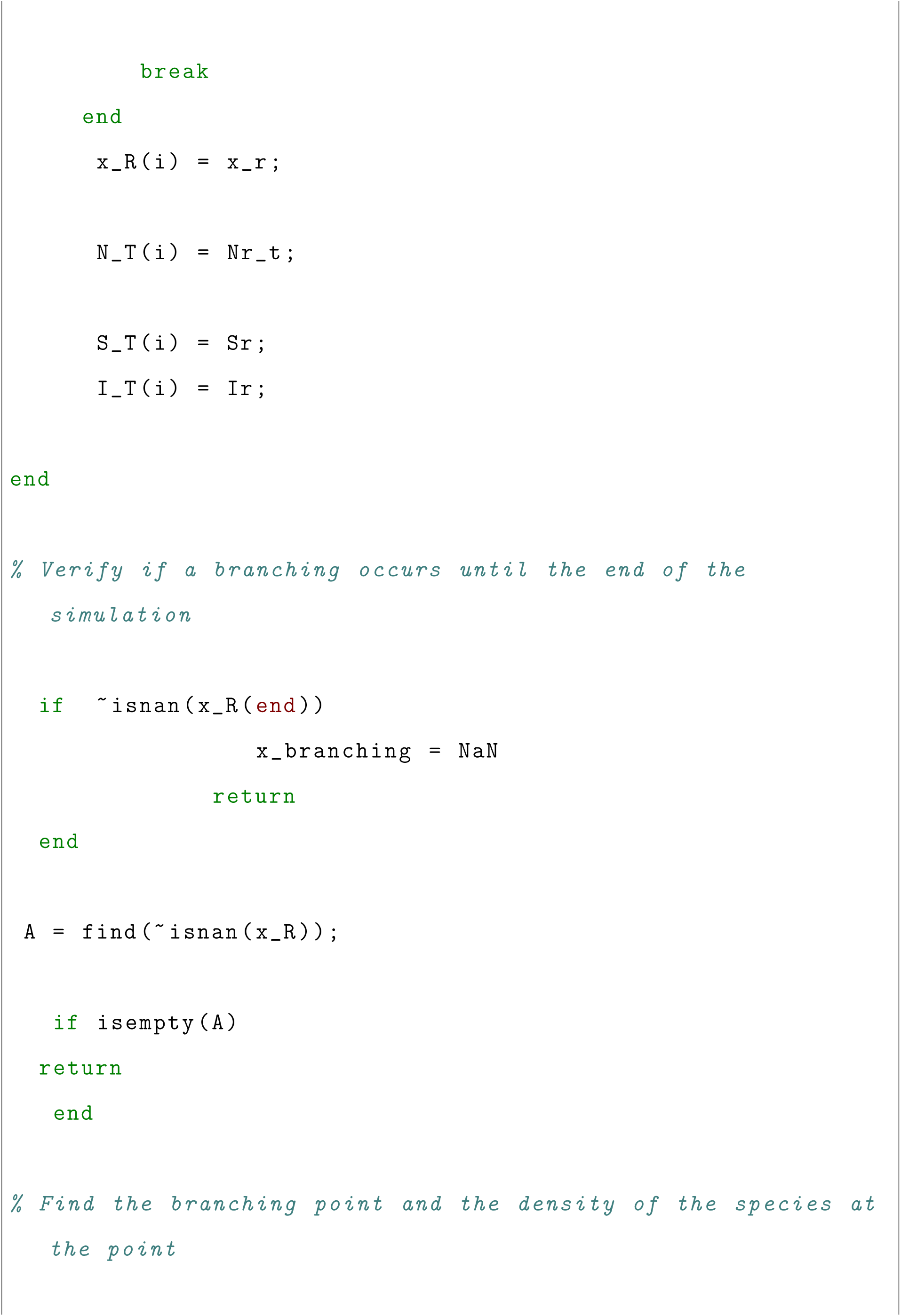
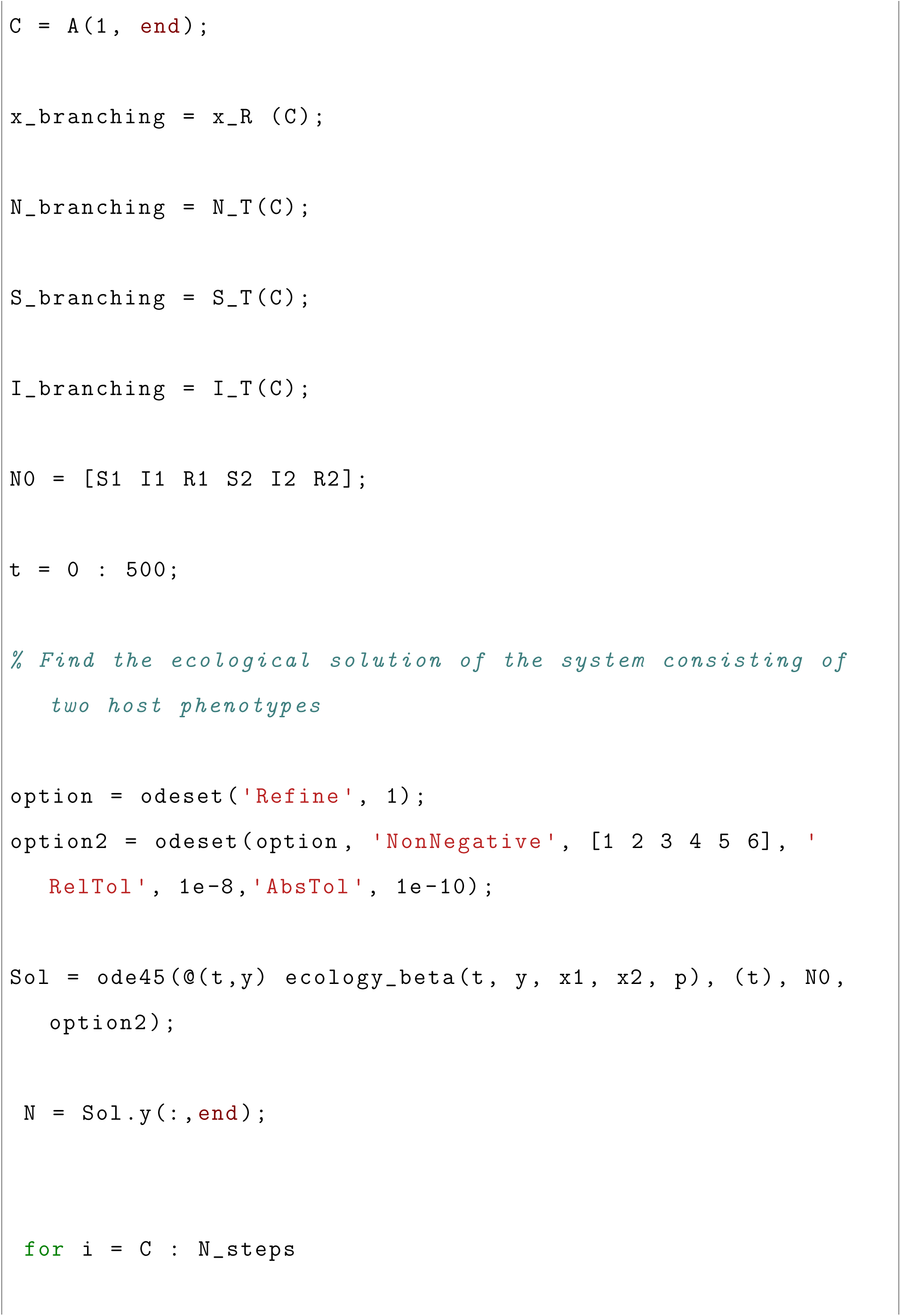
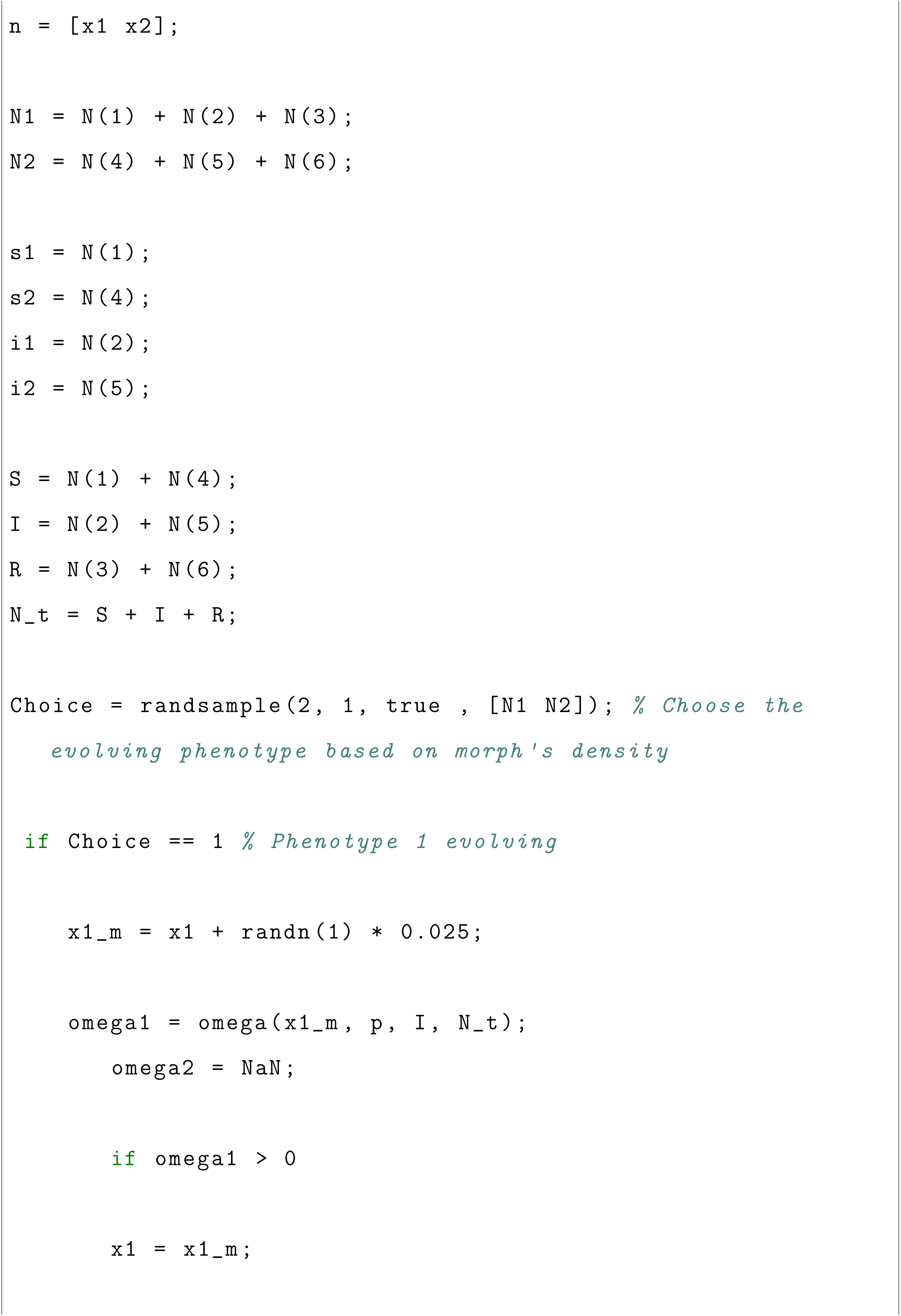
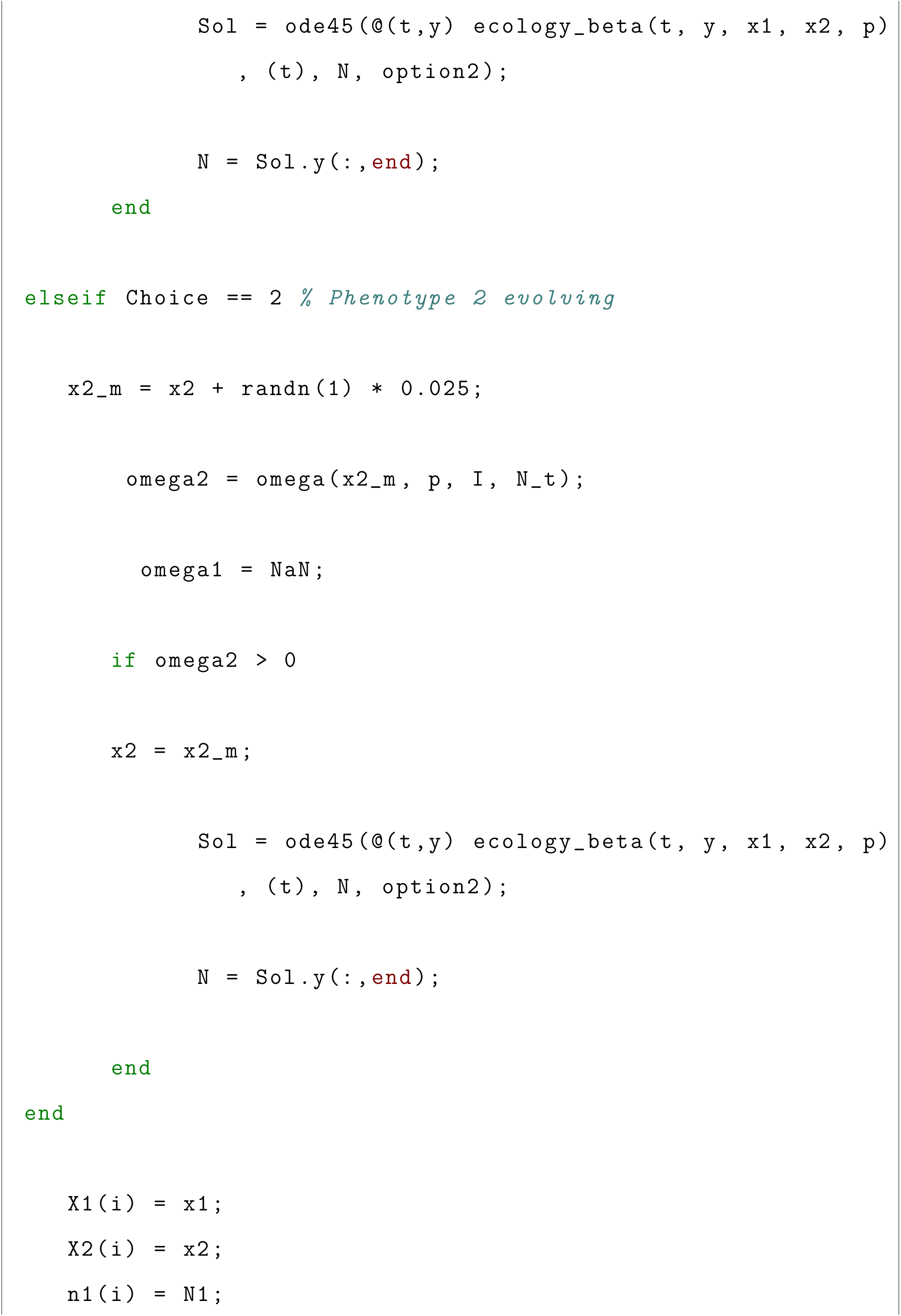
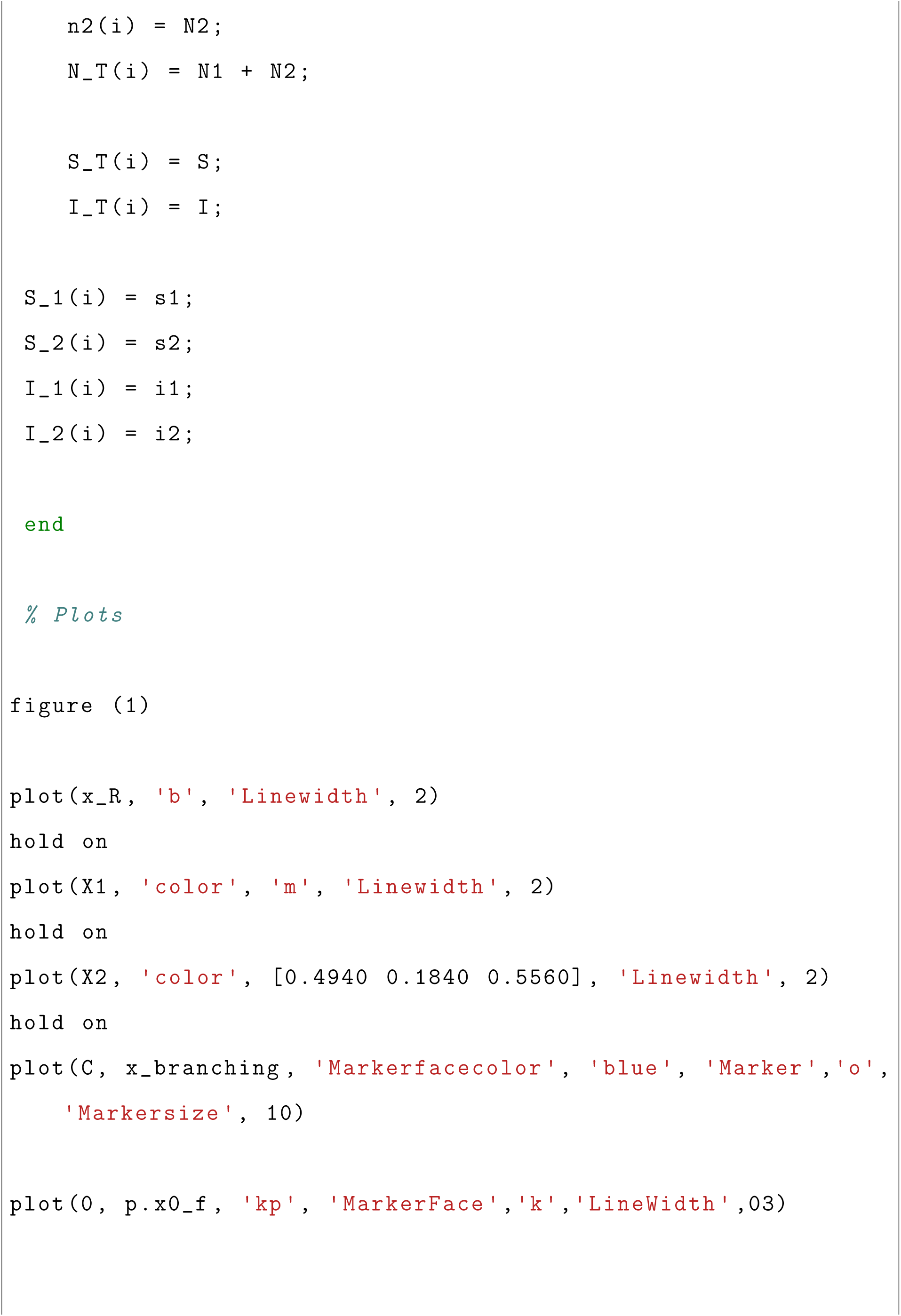
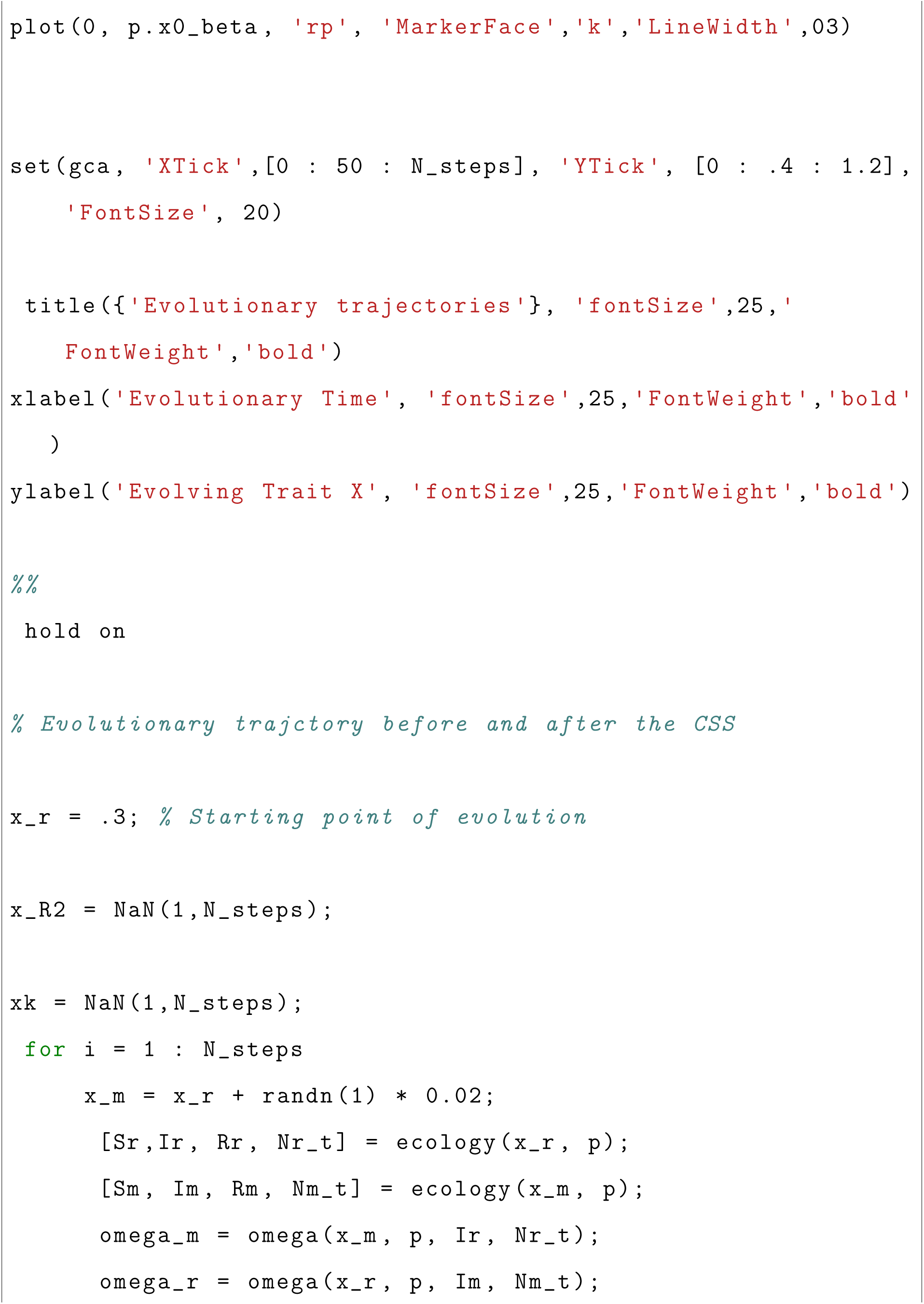
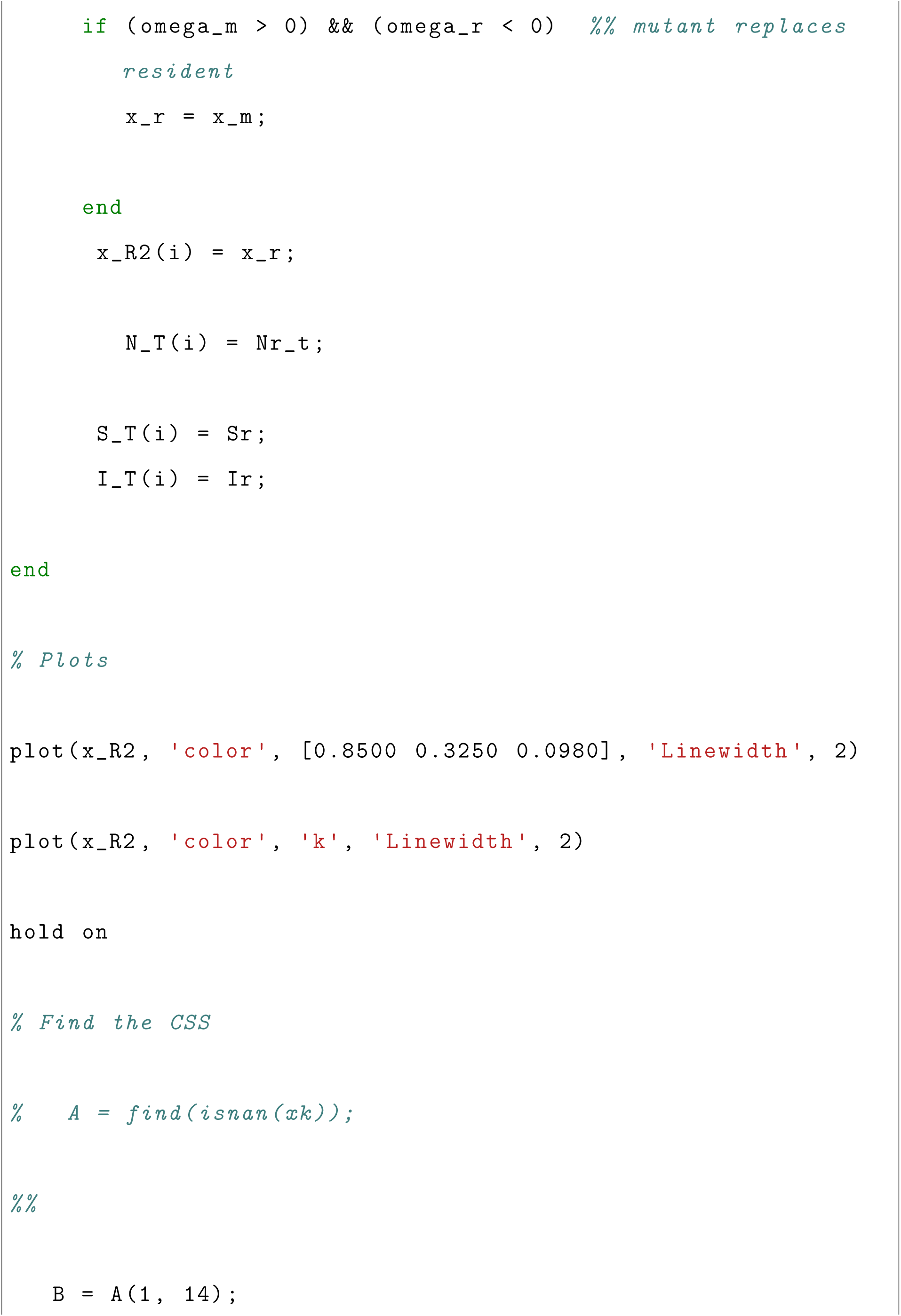
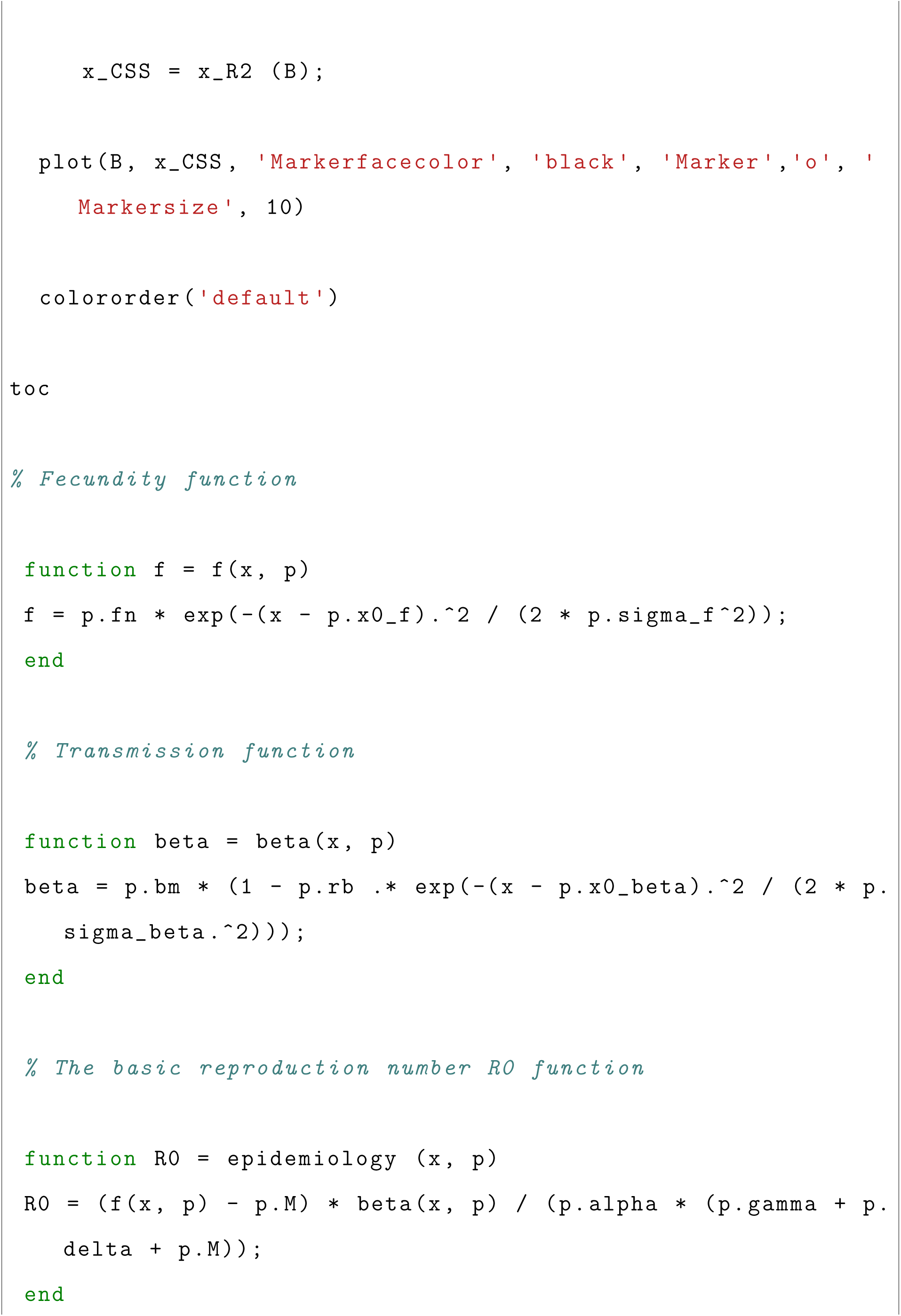
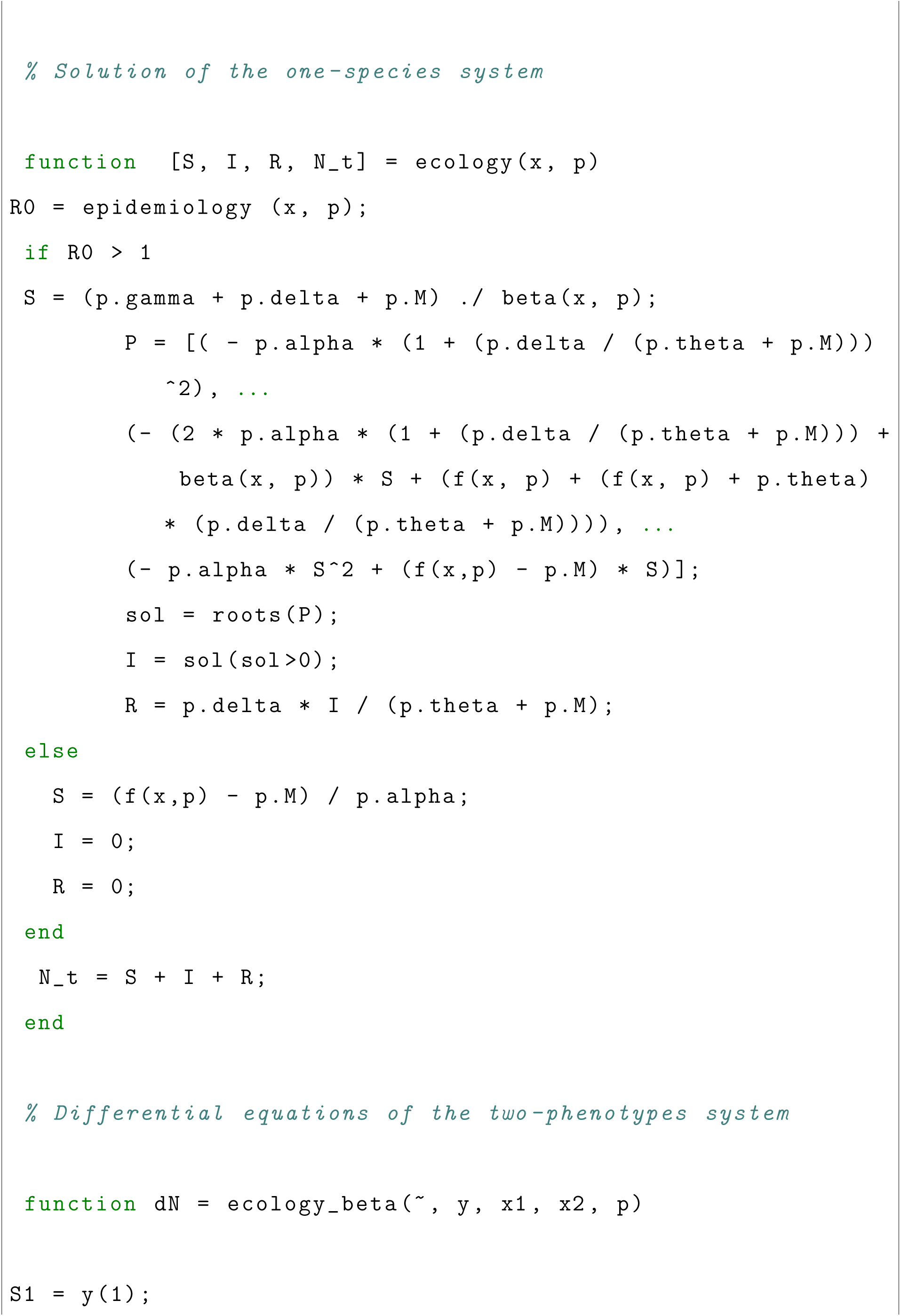
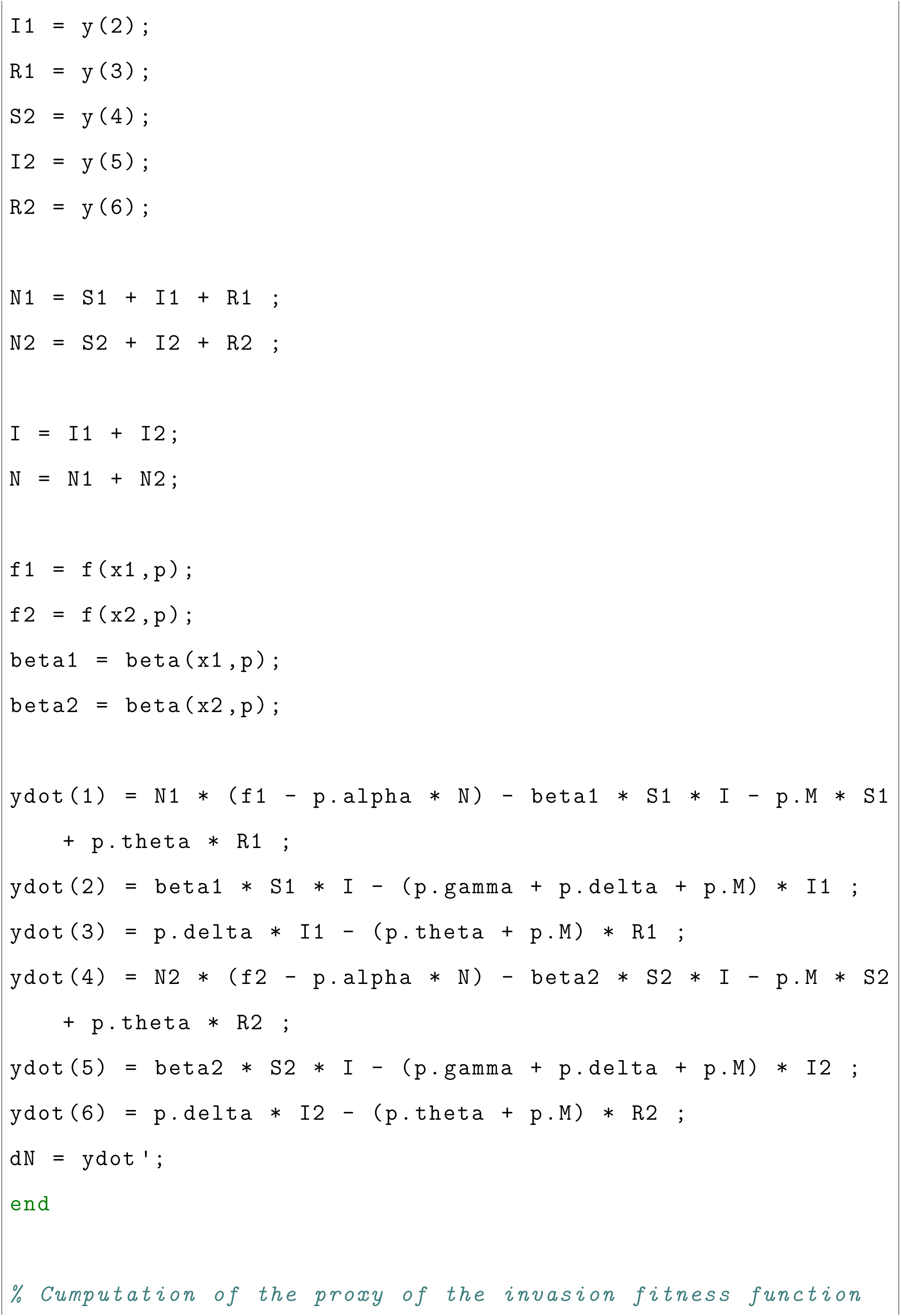
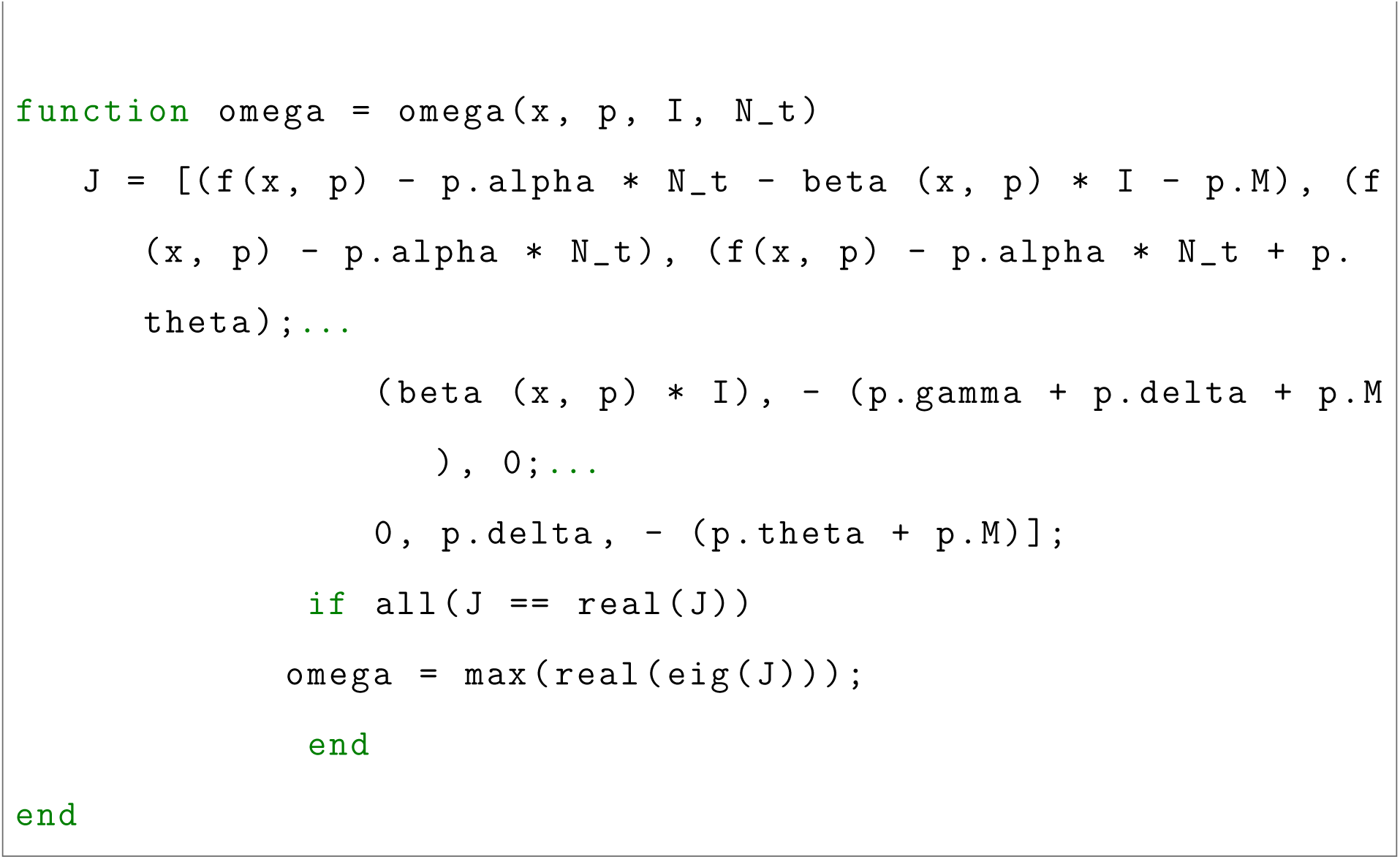

Figure 5.A, D and E, and figure S1

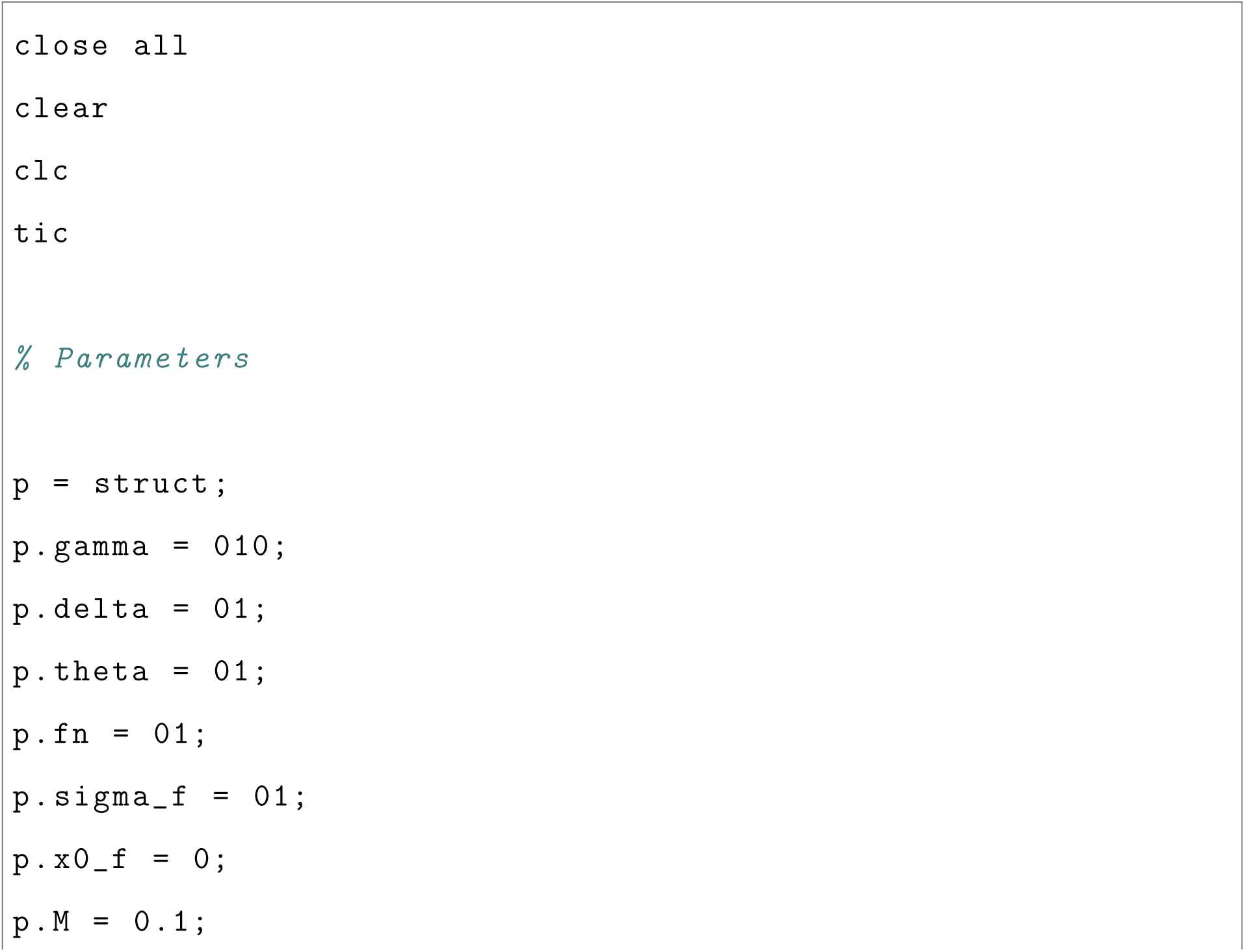
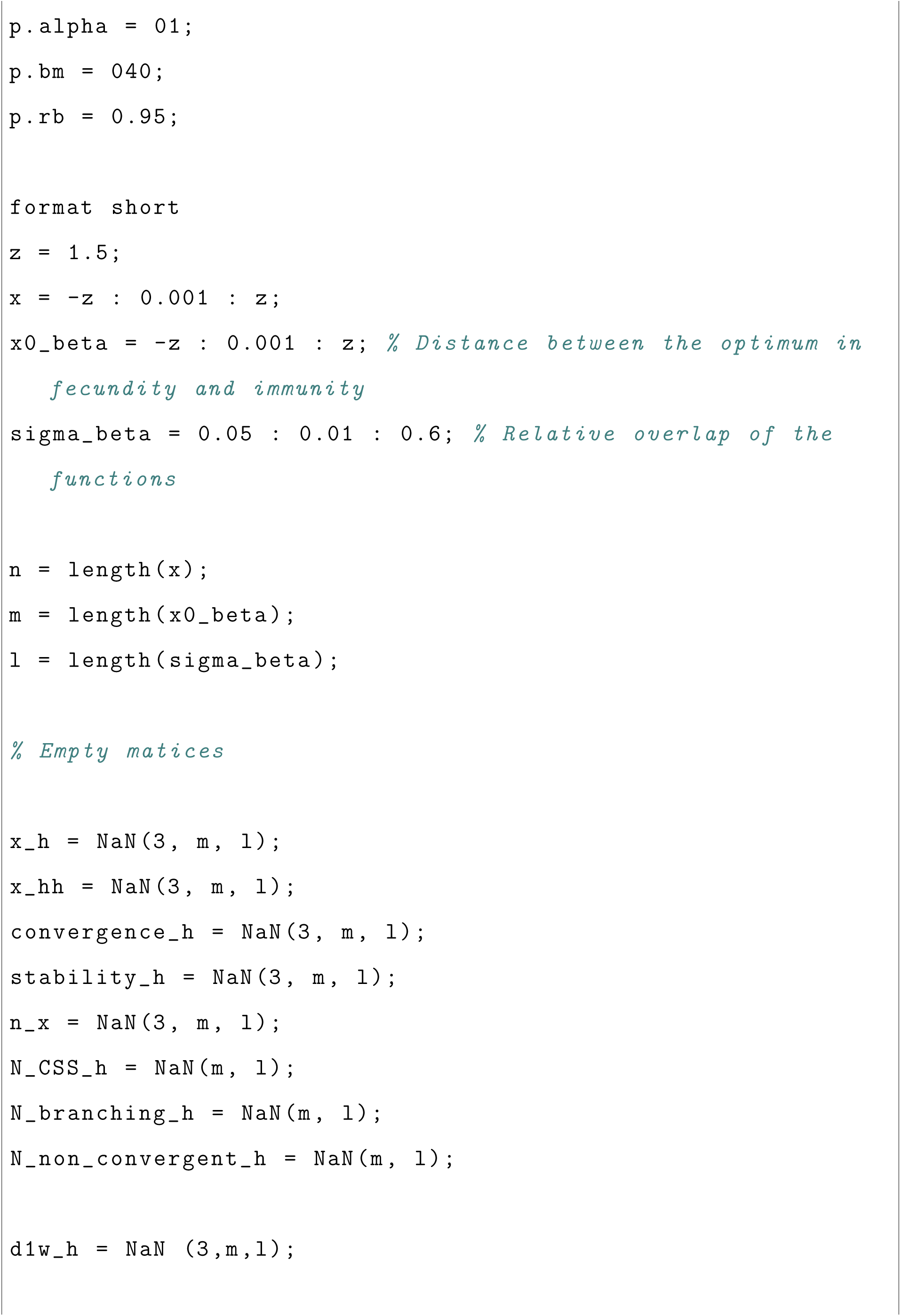
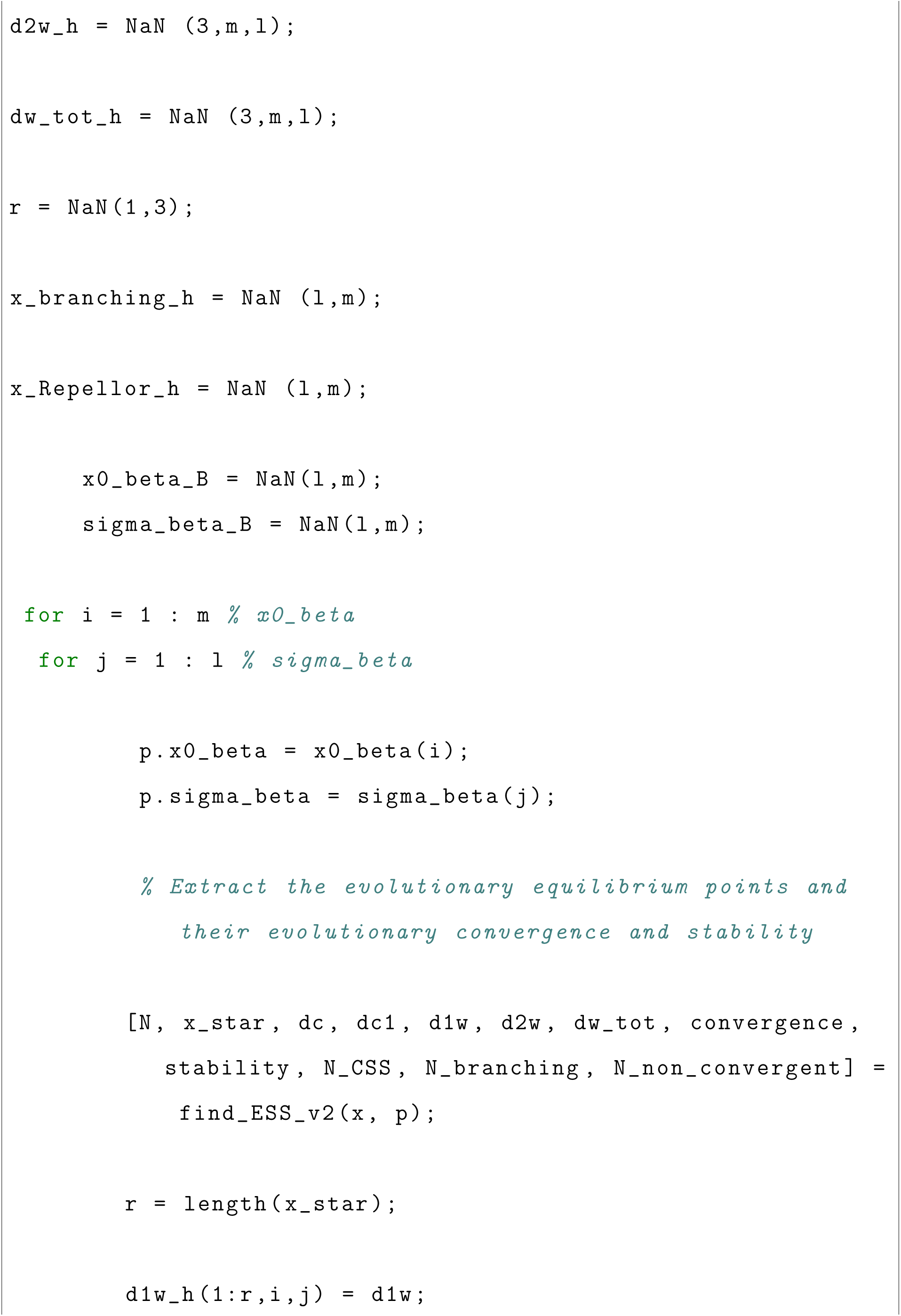
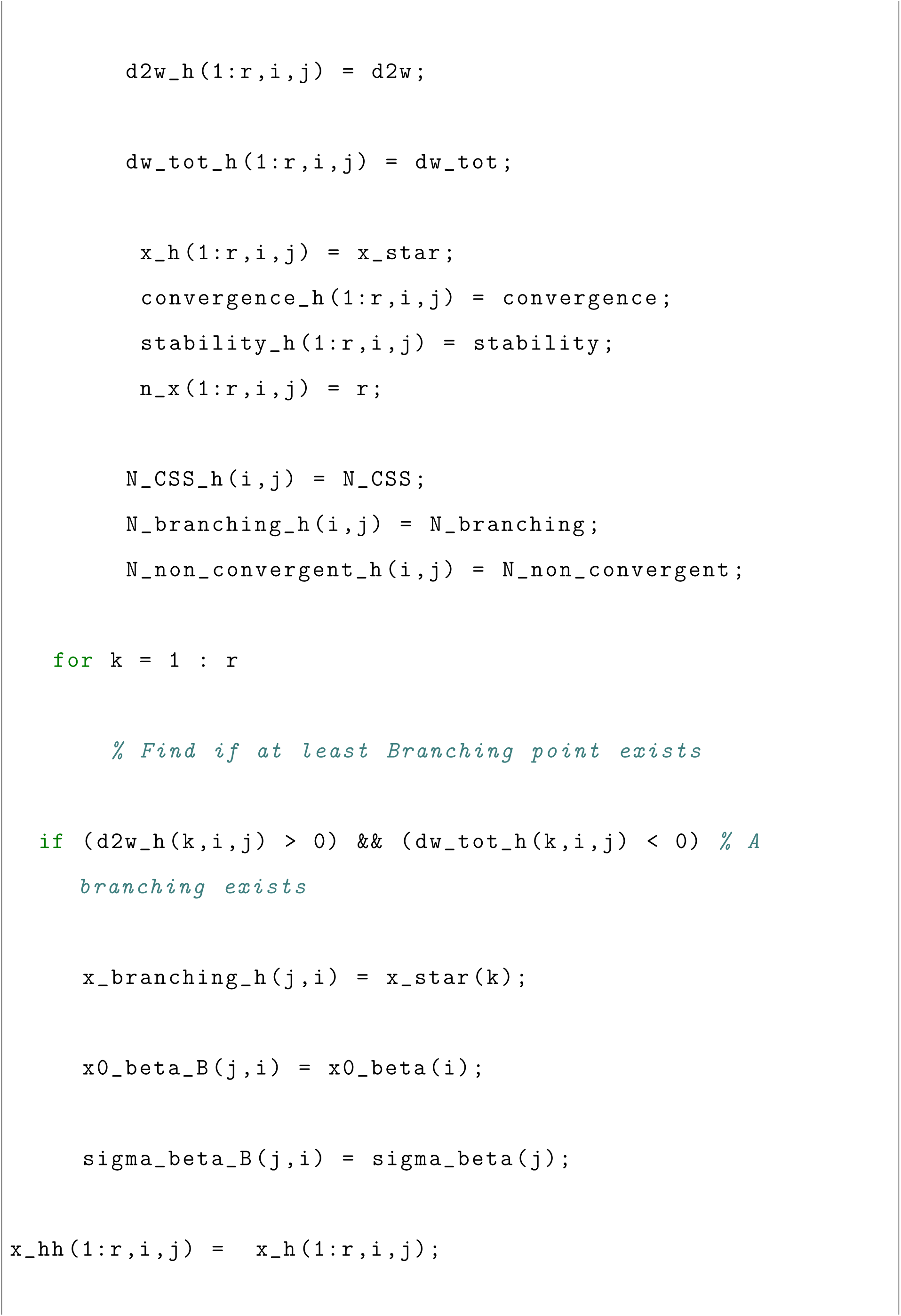
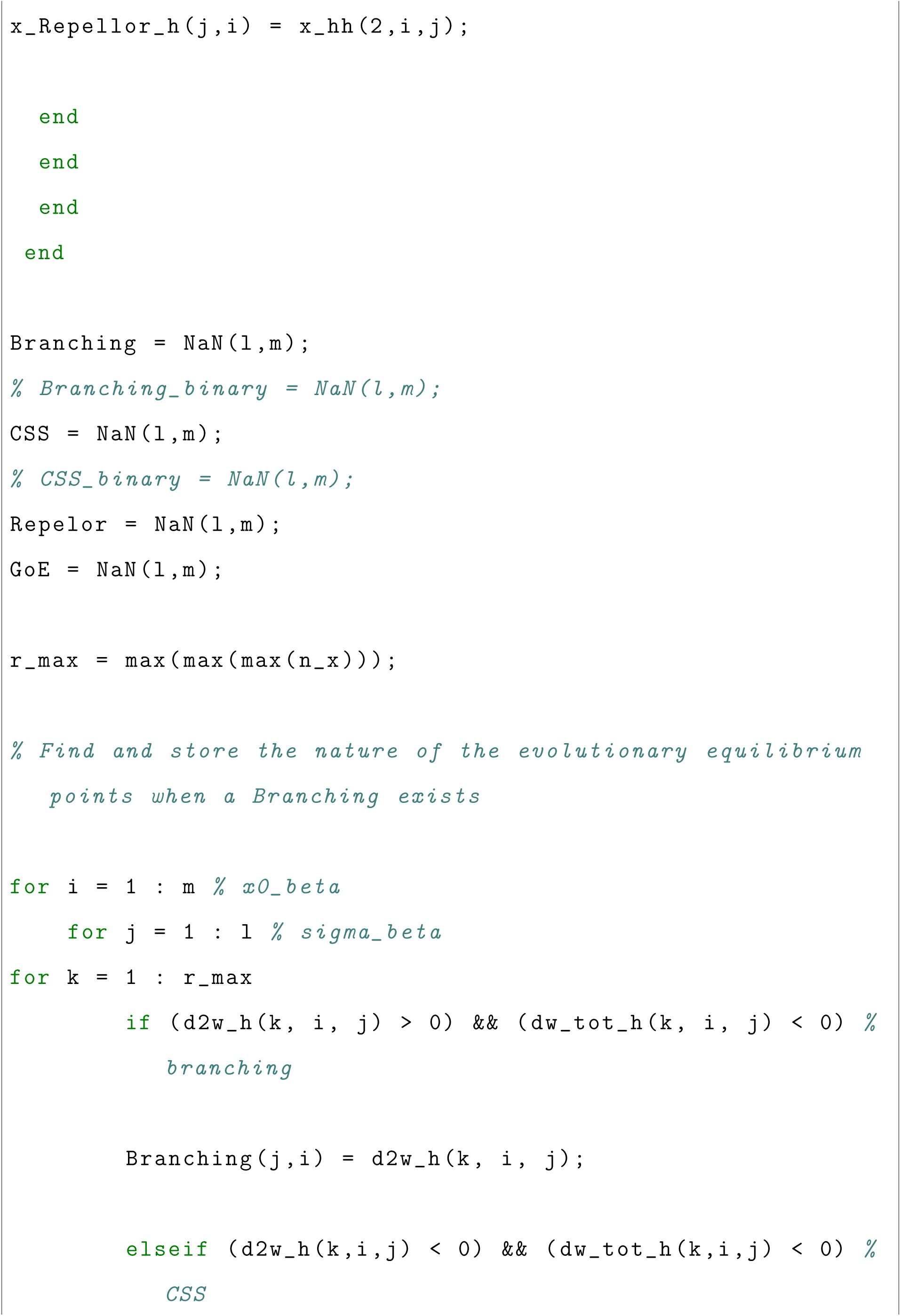
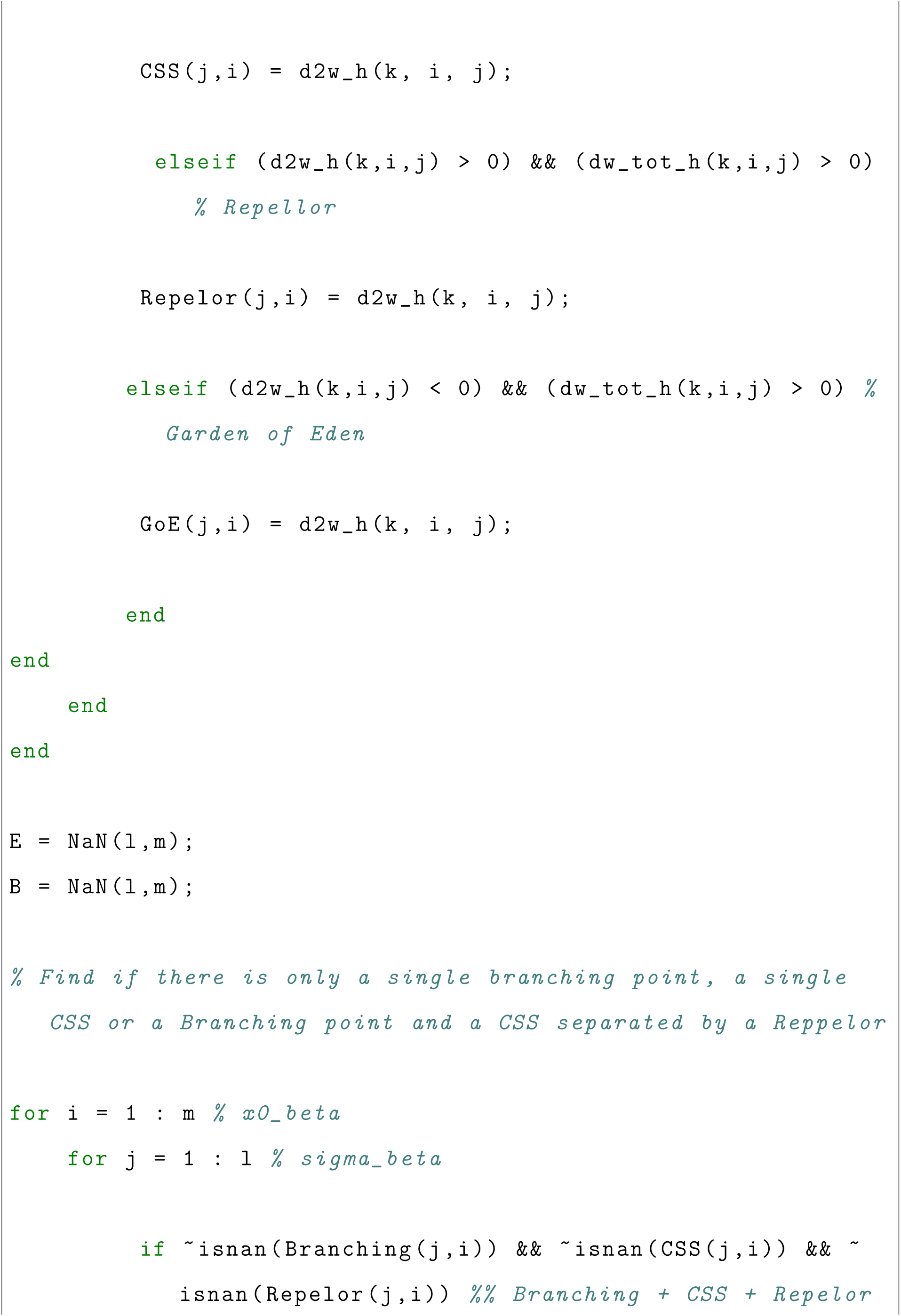
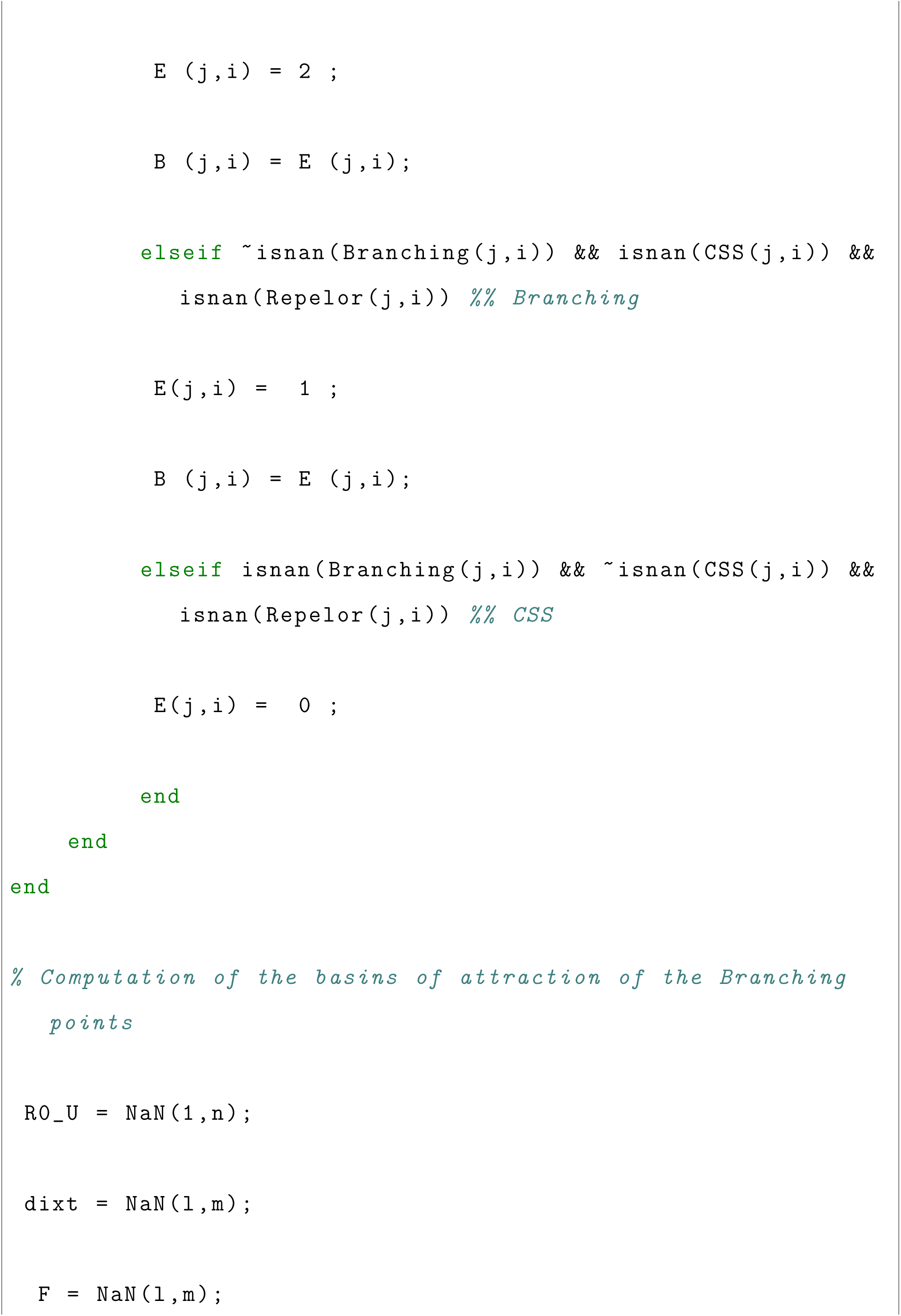
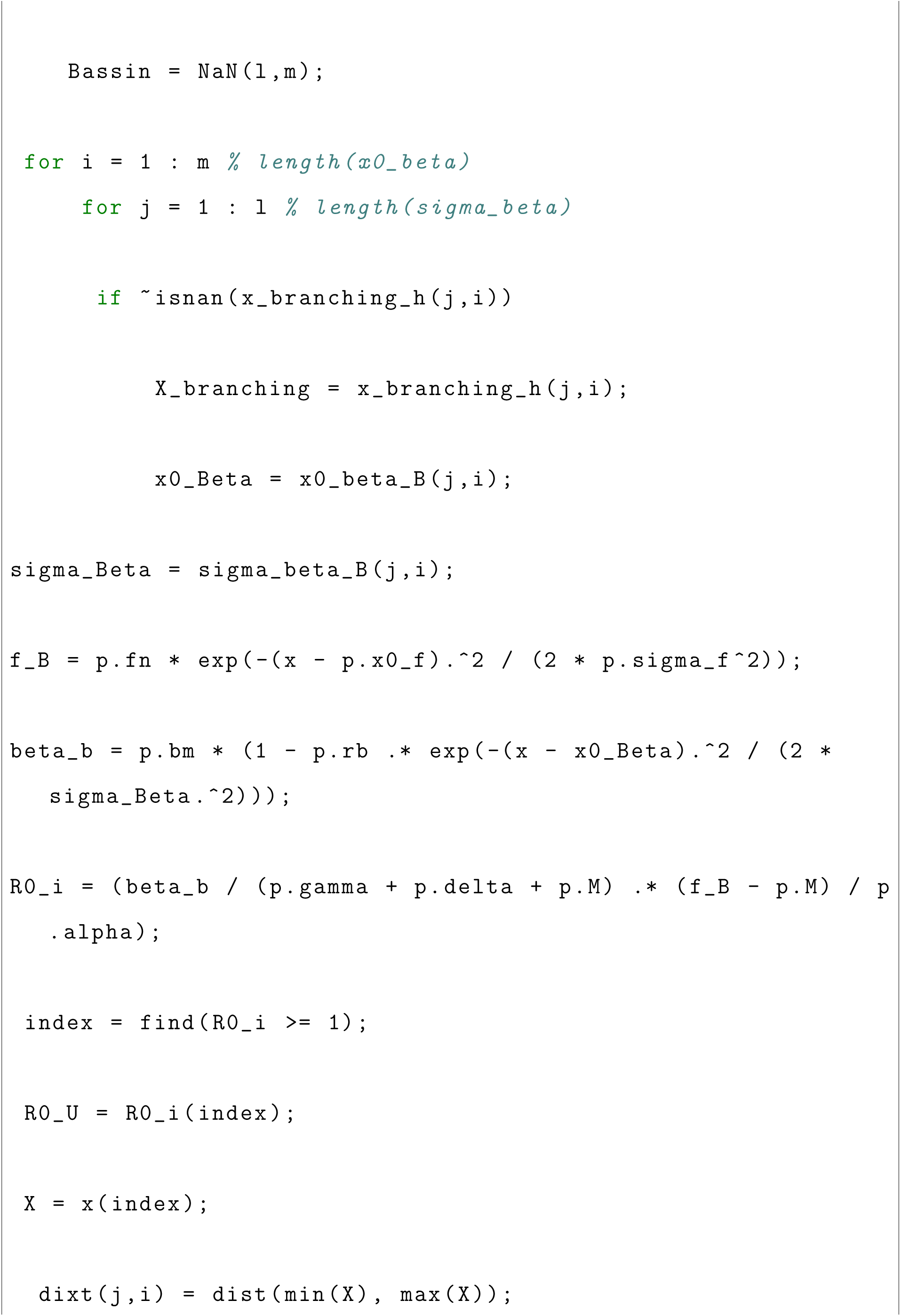
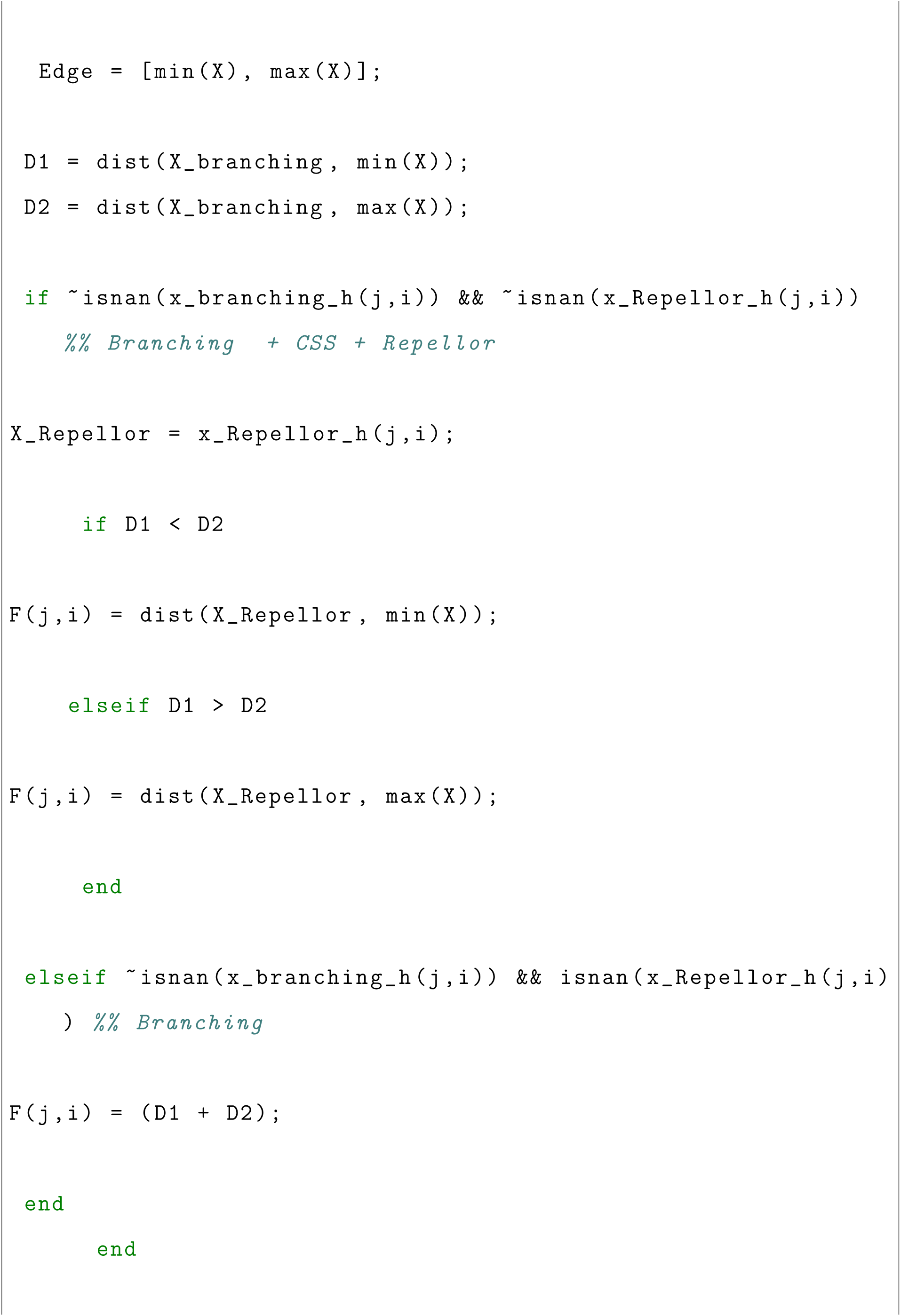
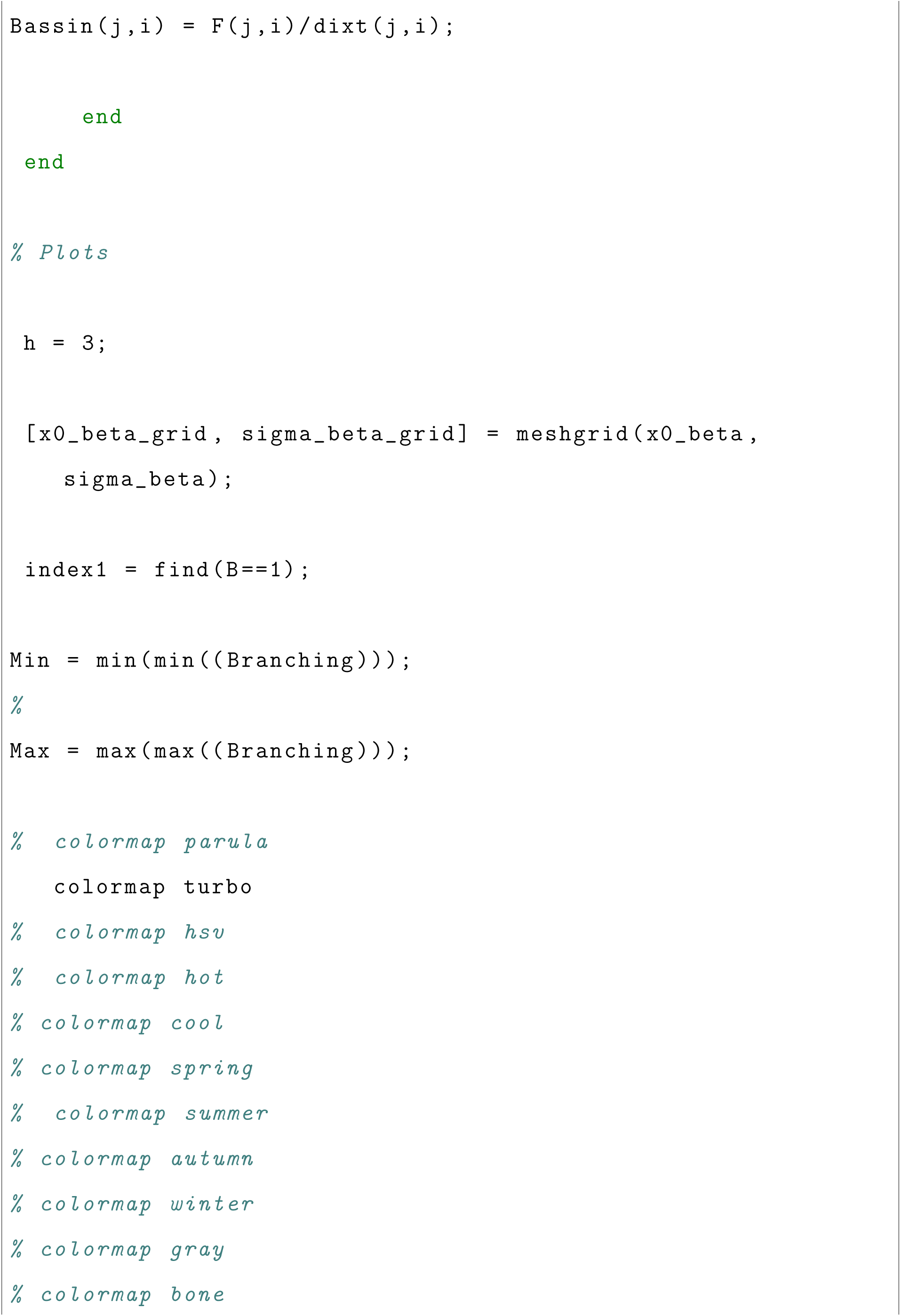
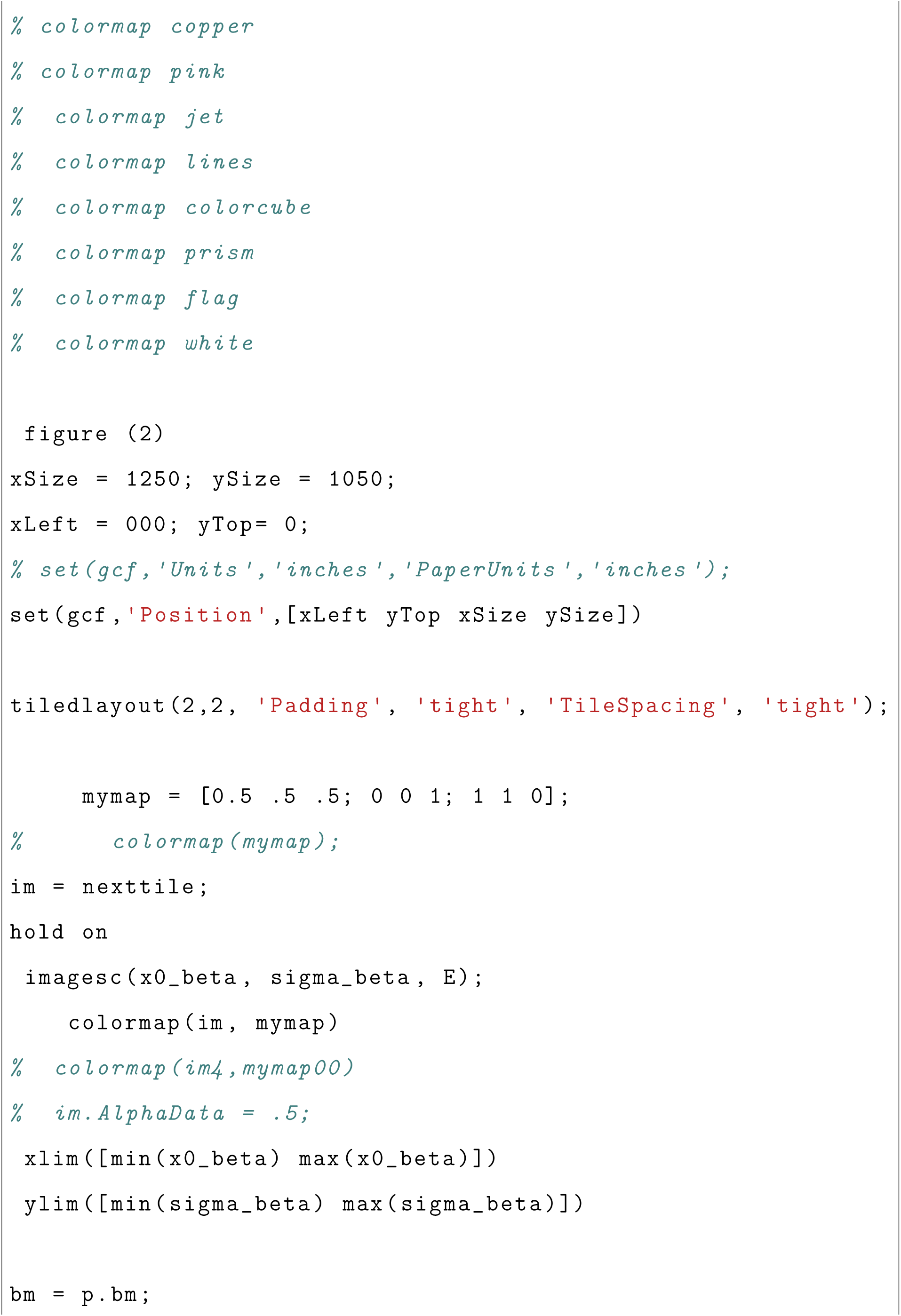
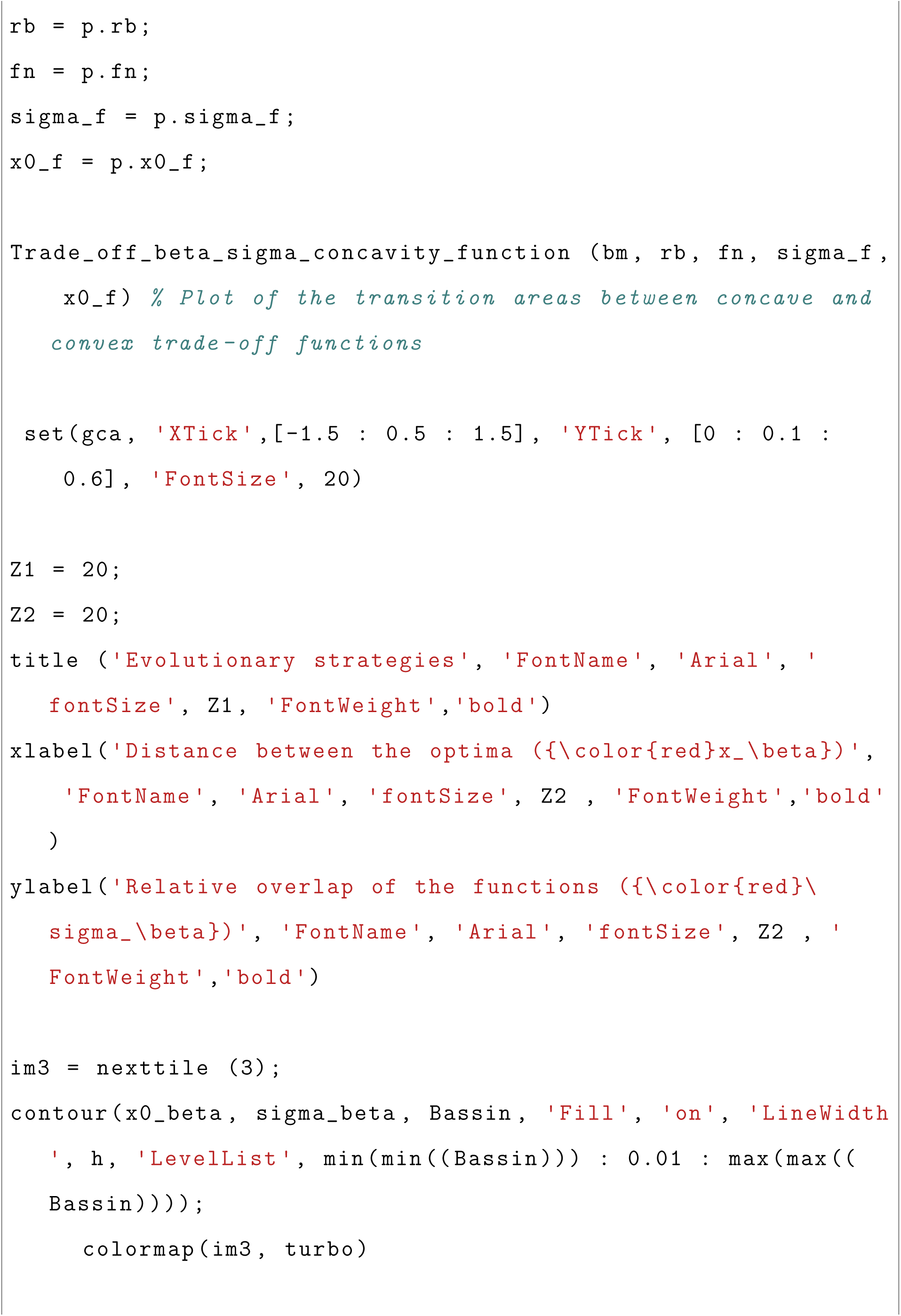
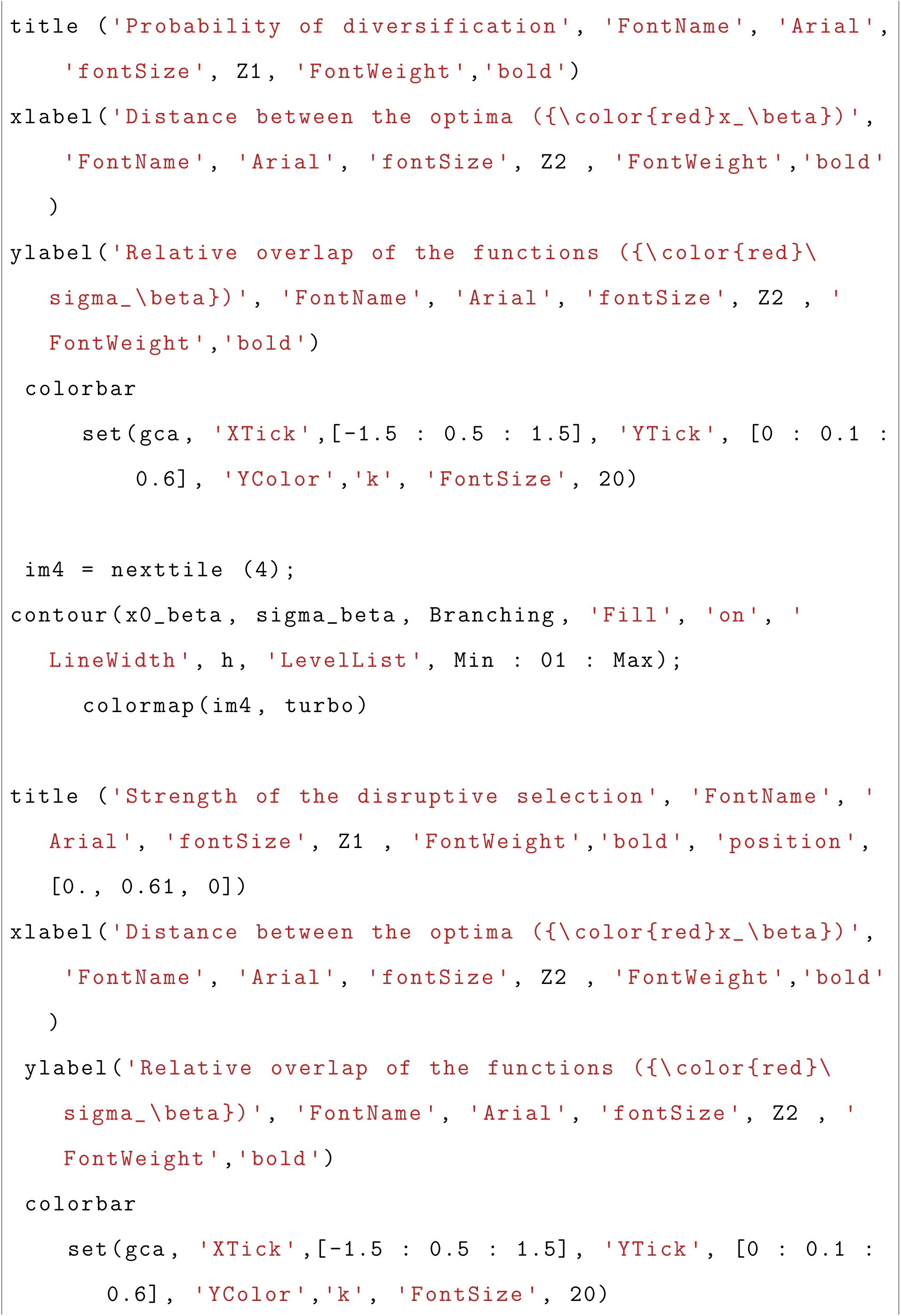
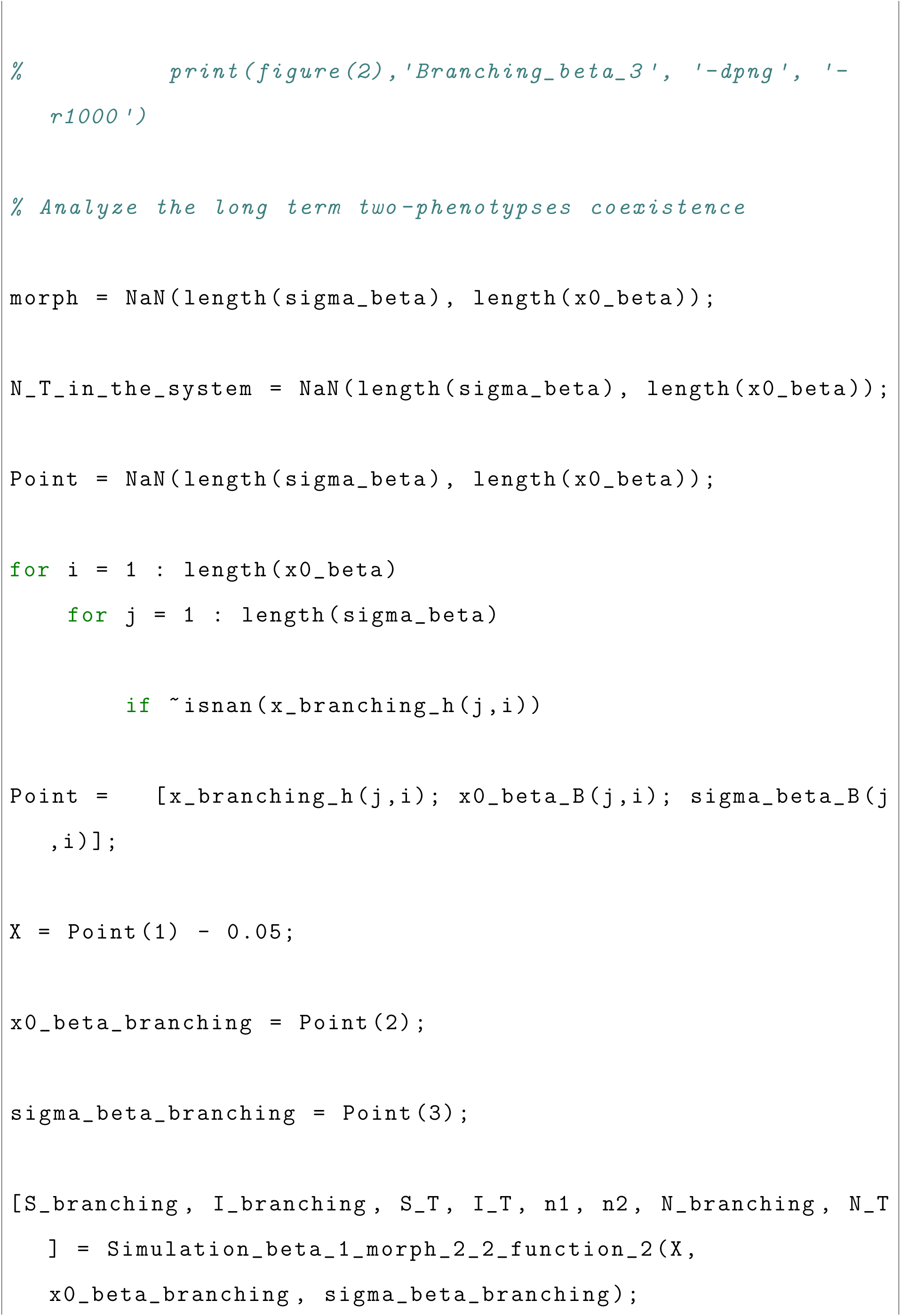
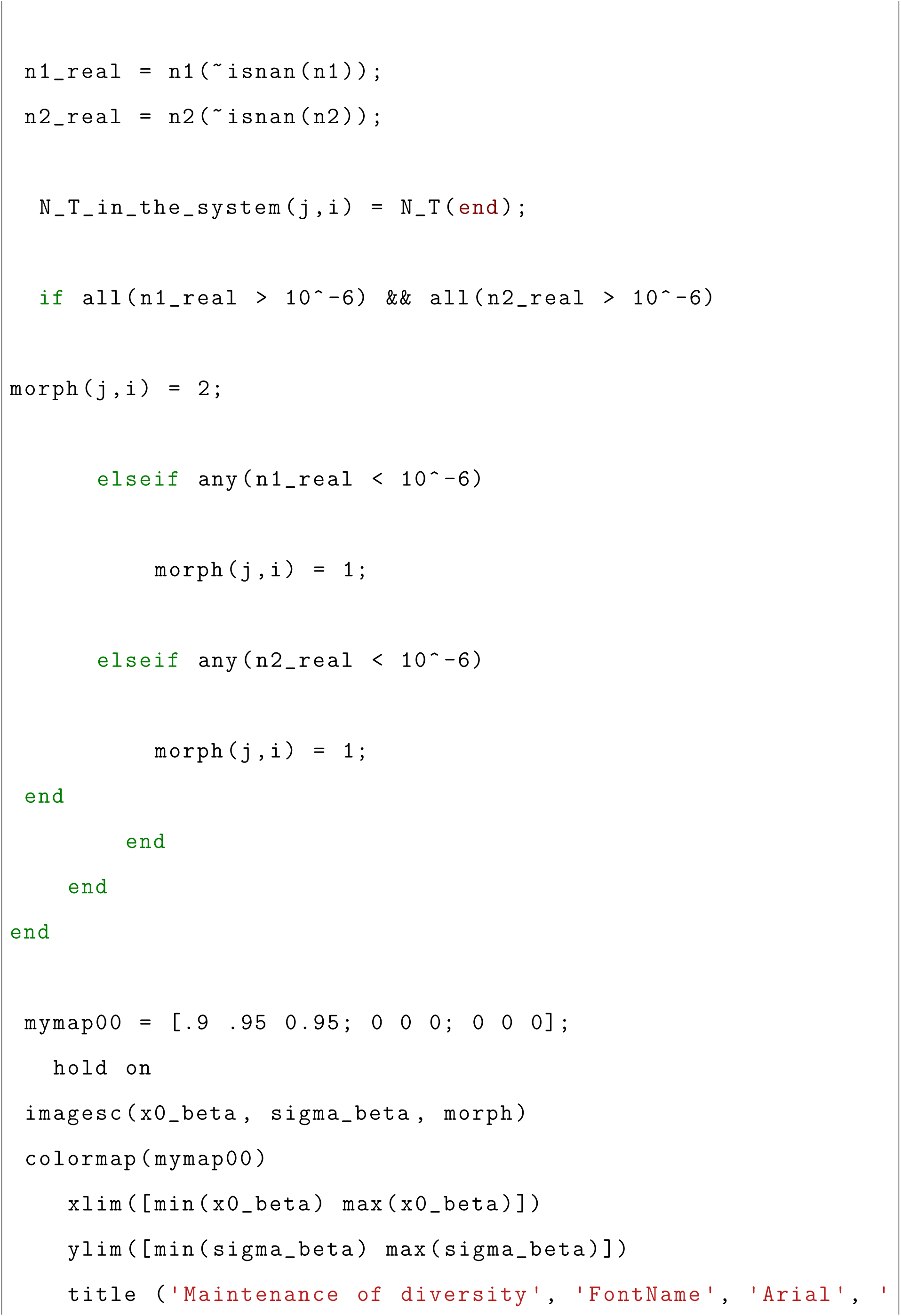
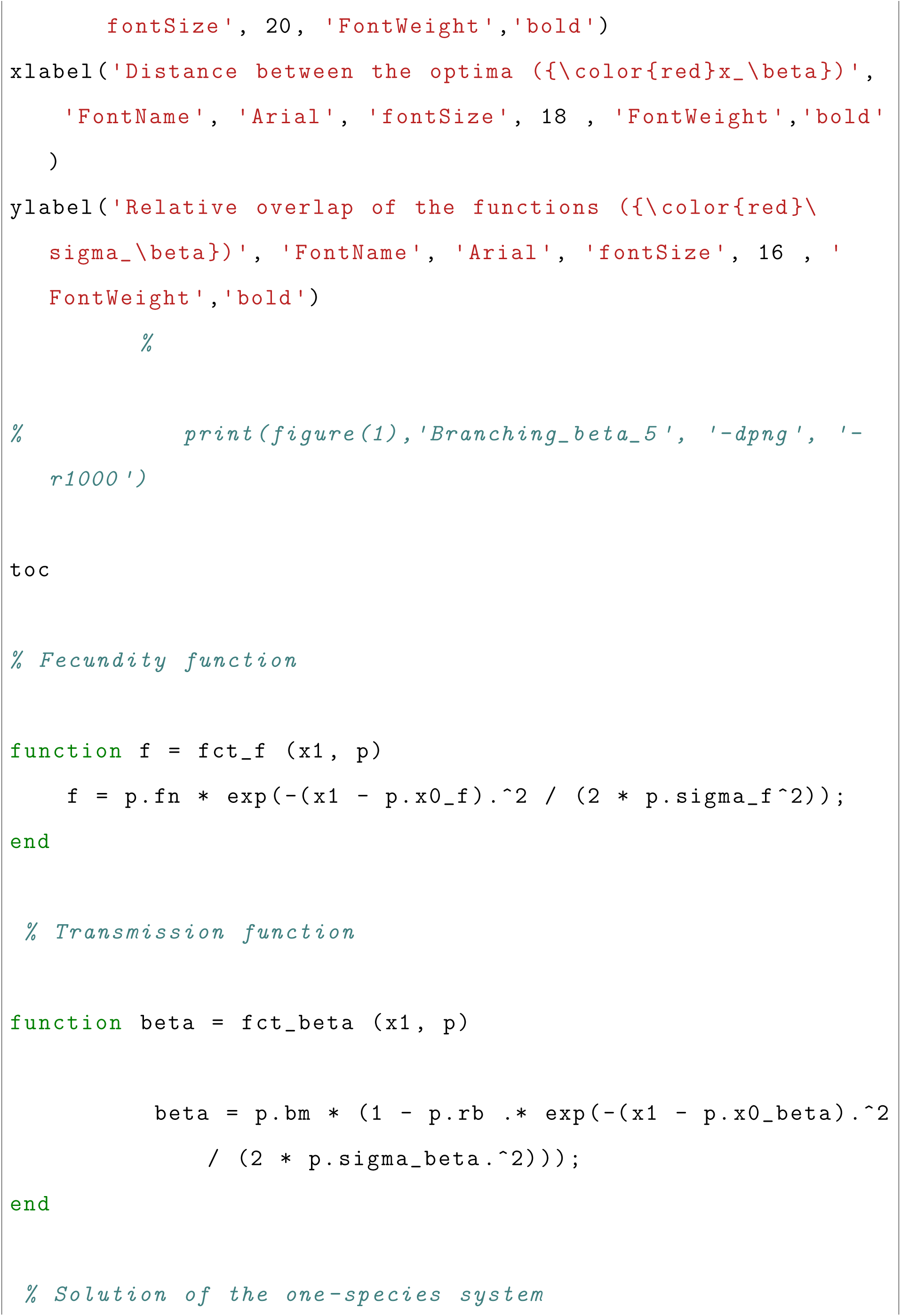
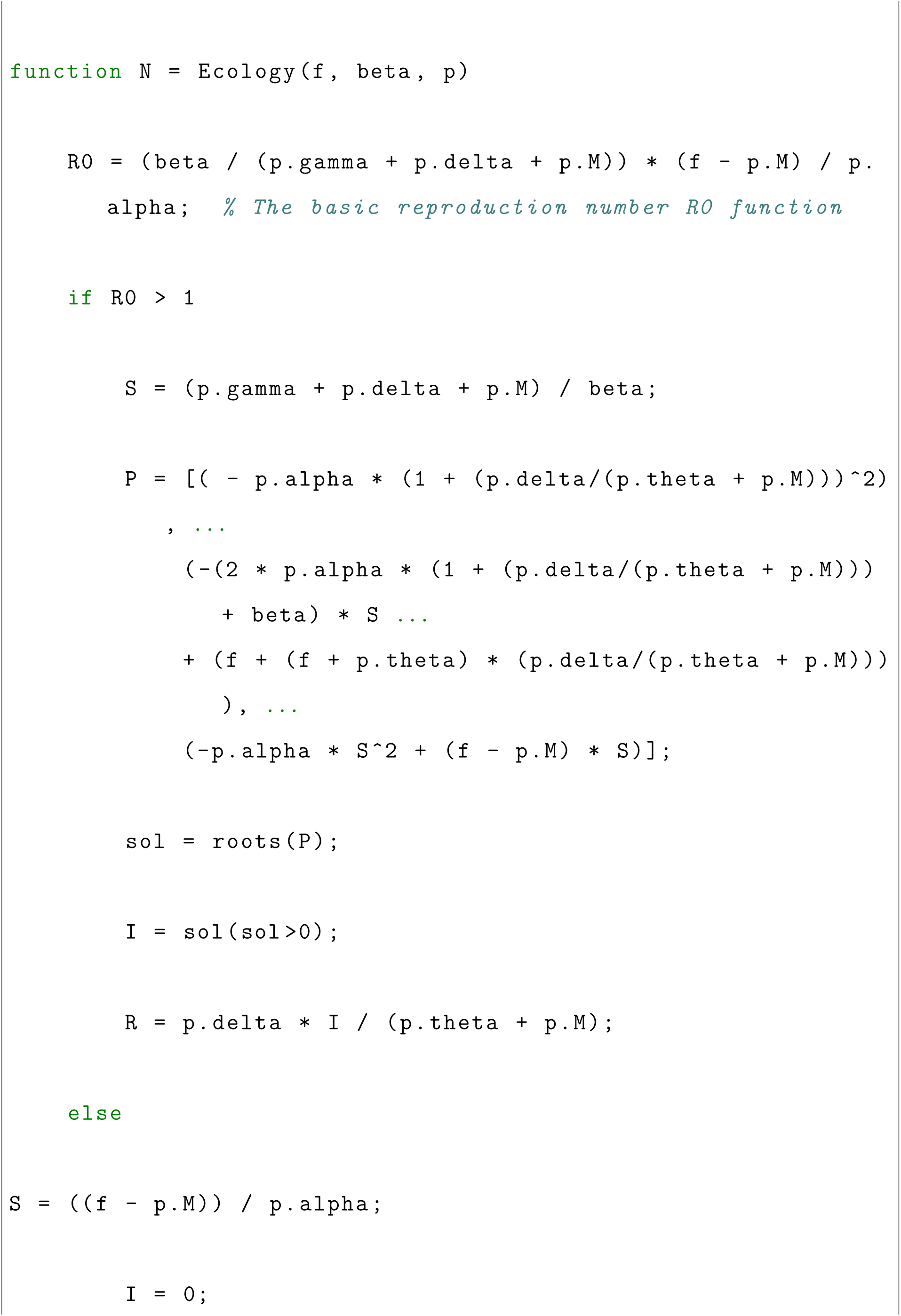
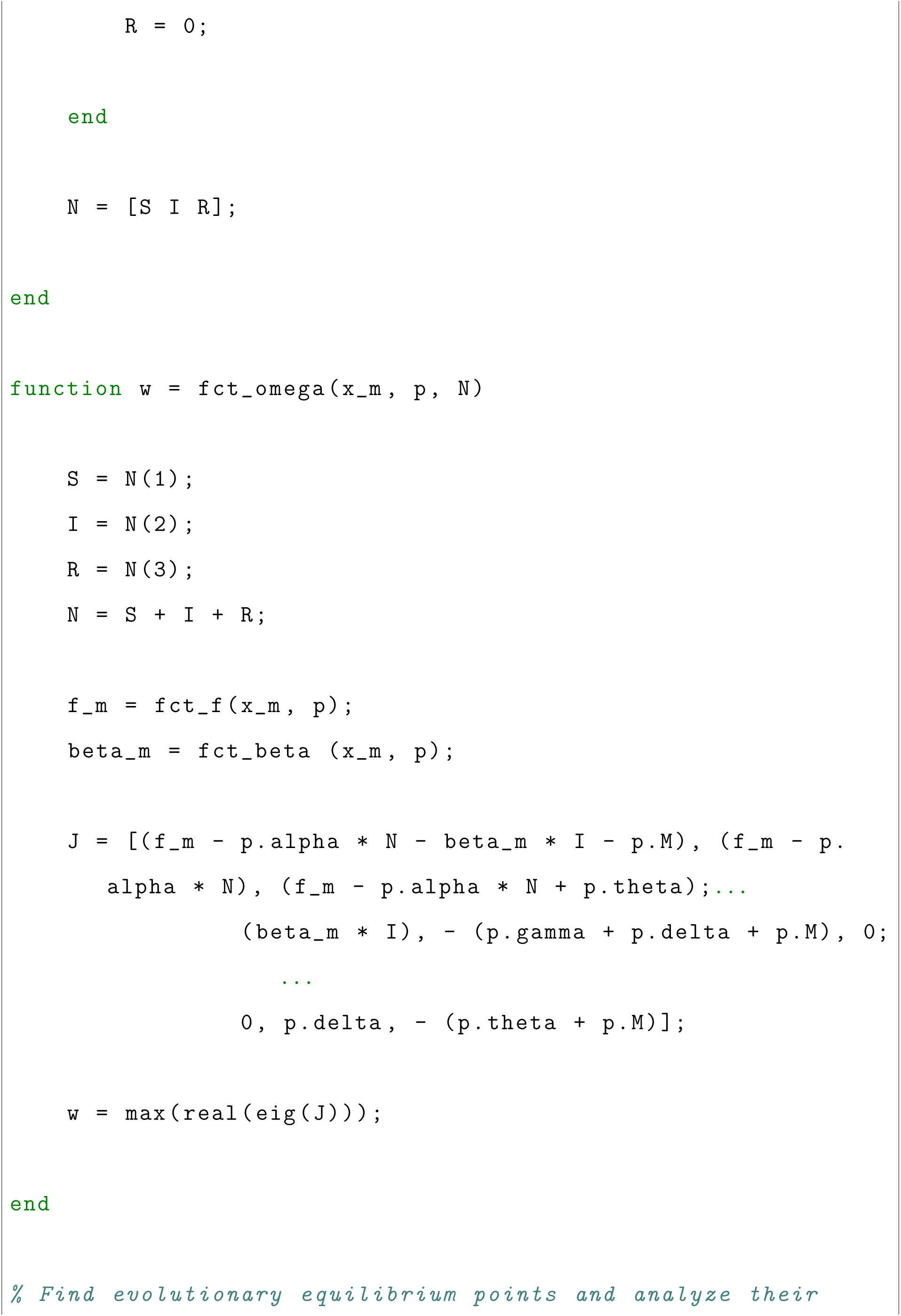
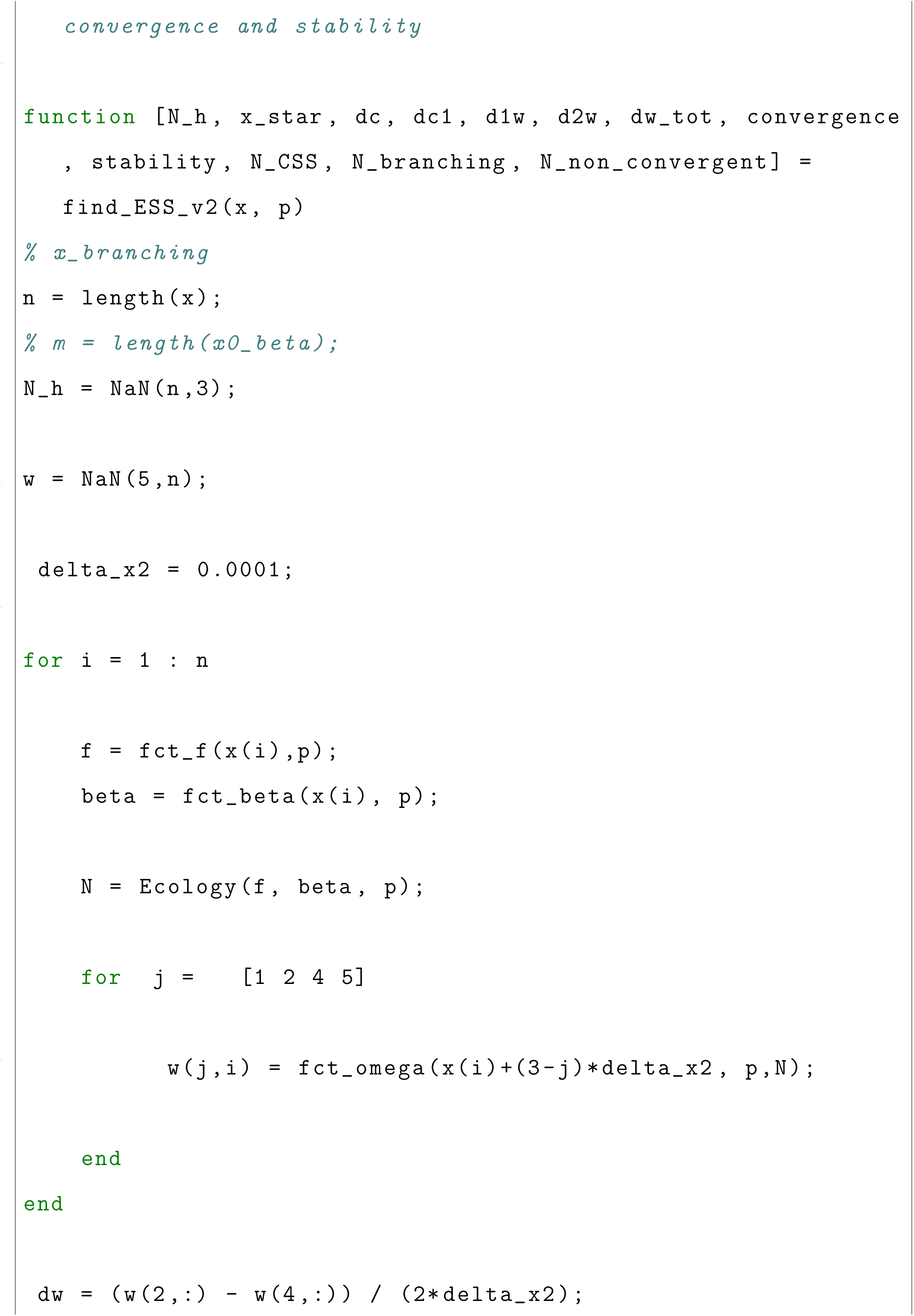
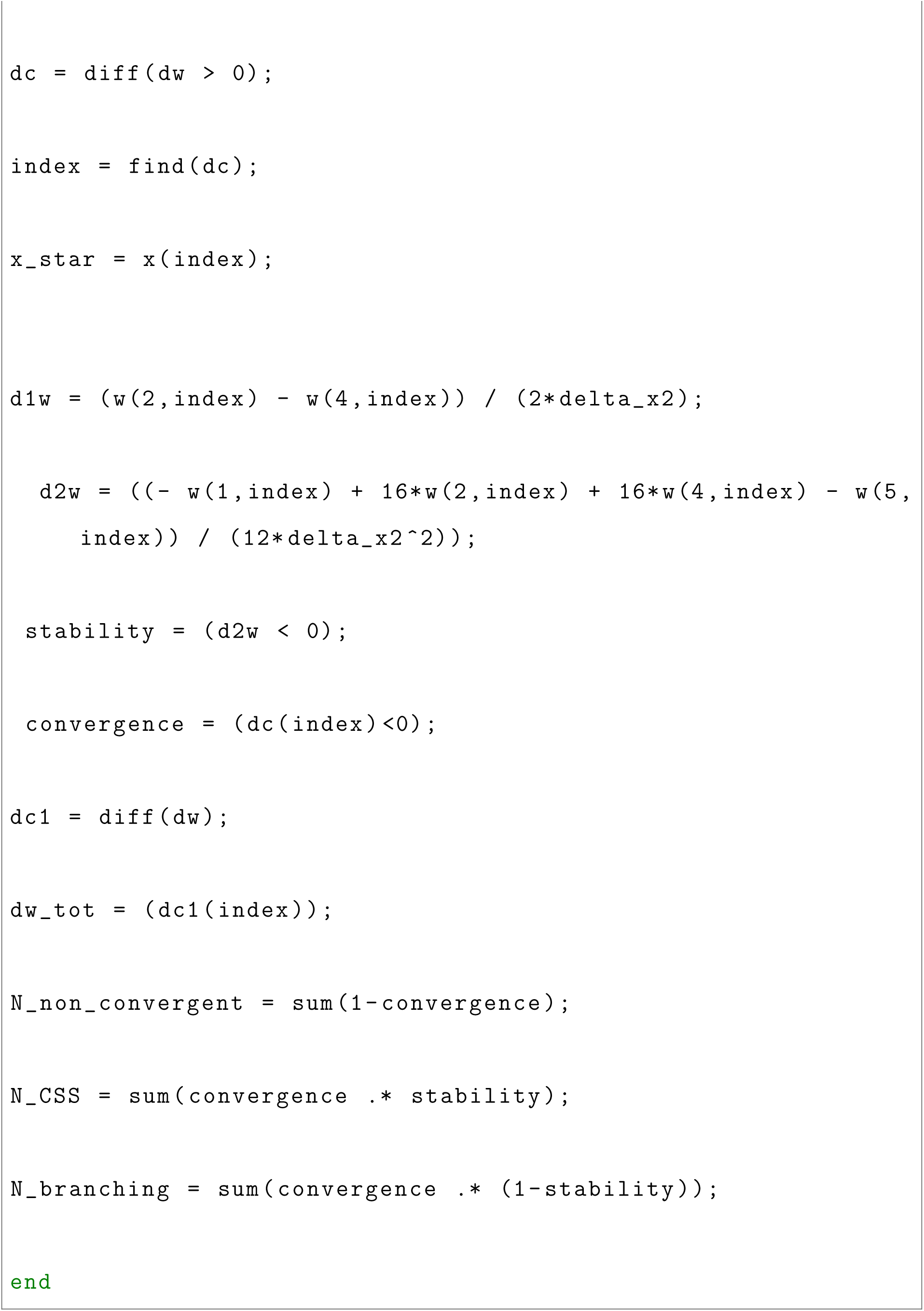

